# Differential Dynamic Behavior of Prefusion Spike Proteins of SARS Coronaviruses 1 and 2

**DOI:** 10.1101/2020.12.25.424008

**Authors:** Vivek Govind Kumar, Dylan S Ogden, Ugochi H Isu, Adithya Polasa, James Losey, Mahmoud Moradi

## Abstract

The coronavirus spike protein, which binds to the same human receptor in both SARS-CoV-1 and 2, has been implied to be a potential source of their differential transmissibility. However, the mechanistic details of spike protein binding to its human receptor remain elusive at the molecular level. Here, we have used an extensive set of unbiased and biased microsecond-level all-atom molecular dynamics (MD) simulations of SARS-CoV-1 and 2 spike proteins to determine the differential dynamic behavior of prefusion spike protein structure in the two viruses. Our results indicate that the active form of the SARS-CoV-2 spike protein is more stable than that of SARS-CoV-1 and the energy barrier associated with the activation is higher in SARS-CoV-2. Our results also suggest that not only the receptor binding domain (RBD) but also other domains such as the N-terminal domain (NTD) could play a role in the differential binding behavior of SARS-CoV-1 and 2 spike proteins.

## Introduction

The etiological agent for the coronavirus disease 2019 (COVID-19) pandemic is SARS-CoV-2, a lineage B *Betacoronavirus* that originated in China towards the end of 2019 ^1–5^. This coronavirus has continued to spread across the world, with millions of confirmed cases and over a million deaths only within a year. Studies have shown that SARS-CoV-2 is more easily transmissible between humans in comparison to SARS-CoV-1 ^6–9^, another lineage B *Betacoronavirus* that caused the 2003 severe acute respiratory syndrome (SARS) epidemic ^10–12^. The differential transmissibility of SARS-CoV-1 and 2 may partially explain the difference in the scale of the SARS epidemic and the COVID-19 pandemic. However, given the striking similarity of the two viruses, the molecular-level explanation of their differential transmissibility is largely missing and has an important implication in developing effective therapeutic agents and vaccines for COVID-19 with long-term efficacy.

SARS-CoV-2 shares several highly conserved structural and functional features with SARS-CoV-1 ^1, 13, 14^. The homotrimeric spike protein is possibly the most important of these and plays a definitive role in the viral infection process by mediating recognition of the host cell receptors ^13, 15–17^. For this reason, the spike protein of SARS-CoV-2 is the primary target of several ongoing structure-based drug and vaccine design studies ^18–23^. Several vaccines have recently been approved for use in different countries, including the mRNA-based Pfizer and Moderna vaccines ^24–26^. SARS-CoV-1 (CoV-1) and SARS-CoV-2 (CoV-2) spike proteins have a high sequence identity of approximately 79% ^1^. The RBDs of both spike proteins interact with the human angiotensin-converting enzyme 2 (ACE2) receptor in order to commence the host-cell infection process ^6, 16, 17, 27–31^. Studies have shown that several regions of the CoV-2 spike protein are susceptible to mutations, with the RBD being particularly vulnerable in this regard ^32–35^. It is possible that therapeutic agents targeting only the RBD-ACE2 interaction might eventually be rendered in-effective due to the appearance of novel mutant strains. Therefore, diversifying the hot spots of the protein being targeted by therapeutics and vaccines is essential in increasing their long-term efficacy. The current study provides a rational framework for such directions by systematically studying the differential behavior of the CoV-1 and CoV-2 spike proteins, highlighting significant regions of the protein that are involved in the activation process, i.e., a large-scale conformational change in the prefusion spike protein, which occurs prior to ACE2 binding.

Recently, several cryogenic electron microscopy (cryo-EM) and computational studies have shed light on the differential receptor binding behavior of the CoV-1 and CoV-2 spike proteins ^6, 17, 27, 36, 37^. The RBD of the spike protein undergoes a large-scale conformational transition from an inactive “down” position to an active “up” position in order to access the ACE2 receptors on the host-cell surface ^6, 17, 27, 38–40^. Experimental studies investigating the binding affinity of the spike protein RBD for the ACE2-peptidase domain (PD) have produced varying results. Using surface plasmon resonance (SPR) and flow cytometry techniques, respectively, Wrapp et al. ^27^ and Tai et. al. ^17^ have reported that the CoV-2 RBD has a higher binding affinity for ACE2-PD than the CoV-1 RBD. For instance, the SPR-based assay shows that the dissociation constant of the CoV-2 spike protein (*K_d_ ≈* 14.7 *nM*) is 10-20 times lower than that of the CoV-1 spike protein ^27, 41^. In a different study, biolayer interferometry has shown that the CoV-2 dissociation constant (*K_d_ ≈* 1.2 *nM*) is only 4 times lower than that of CoV-1, indicating that the binding affinities are generally comparable ^6^. Such quantitative inconsistencies emphasize the need to improve our understanding of the mechanistic aspects of the RBD-ACE2 interaction. A disadvantage of experimental techniques like SPR and biolayer interferometry is that they require the protein to be immobilized prior to measuring the binding affinity ^42, 43^. This introduces a level of bias into these experimental assays, particularly if the binding behavior of a protein is conformation-dependent, as is the case for the coronavirus spike proteins. The imposed protein immobilization thus causes the loss of valuable information regarding the conformational changes that lead to spike protein activation. One may argue that many studies so far have neglected the fact that the binding process involves not only the RBD-ACE2 interaction but also the spike protein activation, a large-scale conformational change with a potentially significant contribution to the differential binding behavior of SARS-CoV-1 and 2. Therefore, to gain a clearer understanding of the enhanced infectivity of SARS-CoV-2, “effective binding” involving both the RBD-ACE2 interaction and the spike protein activation/inactivation process needs to be investigated. Here, we focus on the latter, which has received less attention in the recent literature.

Cryo-EM studies have successfully resolved structures of both spike proteins in the inactive state, active unbound state, and active ACE2-bound state ^6, 27, 31, 38, 44^. However, cryo-EM and X-ray crystallography studies essentially capture static pictures of specific protein conformations and do not provide detailed information on the dynamic behavior that drives major conformational transitions ^45–47^. In addition, given the substantial differences in the experimental and physiological conditions, it is not clear whether all relevant conformational states are captured using techniques such as cryo-EM. For instance, a recent single-molecule fluorescence resonance energy transfer (smFRET) study has captured an alternative inactive conformation for the CoV-2 spike protein ^48^ that is not consistent with those obtained from cryo-EM. It is thus important to investigate the differential conformational landscapes of the CoV-1 and CoV-2 spike proteins in terms of both important functional states and their dynamics. For this purpose, we use an extensive set of microsecond-level unbiased and biased MD simulations. Here, we make certain assumptions to be able to make progress towards deciphering the differential behavior of the two spike proteins, such as relying on cryo-EM structures as our initial models, excluding the unresolved transmembrane domain of the spike protein, and excluding the glycan chains in the simulations. However, we treat the spike proteins of both viruses similarly so that a reliable comparison can be made.

Our extensive all-atom equilibrium MD simulations show that the active CoV-2 spike protein is potentially more stable than the active CoV-1 spike protein. We also report that the RBD of the active CoV-1 spike protein can undergo a spontaneous conformational transition to a pseudo-inactive state characterized by the interaction of the NTD and RBD, a state not observed in any of the previous experimentally reported structures of CoV-1 or CoV-2 spike protein. This observation is broadly in line with the recent smFRET experimental results indicating the potential for the presence of alternative inactive spike protein conformations ^48^. More specifically, electrostatic interaction analyses reveal that unique salt-bridge interactions between the NTD and RBD of the CoV-1 spike protein, are involved in the major conformational transition observed in our simulations. No large-scale conformational changes occur in any of the active CoV-2 spike protein simulations or any of the inactive CoV-1 or CoV-2 spike protein simulations within the timescale of our unbiased MD simulations (5 *µ*s).

In order to investigate the longer timescale conformational dynamics inaccessible to unbiased simulations ^49^, we have also employed extensive steered MD (SMD) simulations. The SMD simulations shed light on the energetics of the conformational change associated with the activation and inactivation processes. The results obtained from these biased simulations strongly suggest that the energy barriers for such conformational transitions, particularly inactivation, are significantly lower for the CoV-1 spike protein and that conformational changes occur more slowly for the CoV-2 spike protein. This provides an explanation for the conformational plasticity displayed by the active CoV-1 spike protein in our simulations as well as the relative conformational stability of the active CoV-2 spike protein. The consistency of results from our equilibrium and nonequilibrium simulations thus provides a reliable picture of the long timescale conformational dynamics of the Cov-1 and CoV-2 spike proteins. The propensity of the active CoV-2 spike protein to maintain the “up” RBD conformation might explain why it has a higher binding affinity for the ACE2 receptor, which in turn could be directly linked to its comparatively high human-to-human transmissibility.

## Results

Recent cryo-EM studies have shown that the prefusion CoV-1 and CoV-2 spike proteins undergo large-scale conformational changes resulting in the exposure of the RBD to the host ACE2 receptors ^6, 38^. However, cryo-EM studies do not provide detailed information on the conformational dynamics of the proteins ^45^. Here, we have used all-atom MD simulations to shed light on the conformational dynamics of the prefusion CoV-1 and CoV-2 spike proteins.

We have performed 5-*µ*s-long unbiased MD simulations of both inactive and active CoV-1 and CoV-2 spike proteins. The active CoV-1 and CoV-2 simulations were repeated additionally twice for another 5 *µ*s each. We have also performed 80 independent nonequilibrium SMD simulations of the CoV-1 and 2 spike proteins, each for 100 ns, to compare the activation and inactivation of CoV-1 and CoV-2 that are otherwise generally inaccessible to unbiased MD. We have thus generated 40 *µ*s of equilibrium and 8 *µ*s of nonequilibrium simulation trajectories in aggregate, results of which are discussed in detail below. All simulations have been performed in an explicit water environment, details of which are discussed in the Methods section.

### The active SARS-CoV-2 spike protein is more stable than the active SARS-CoV-1 spike protein

Within the timescale of our unbiased equilibrium simulations (i.e., 5 *µ*s), the inactive forms of both CoV-1 and CoV-2 spike proteins do not undergo any major conformational transitions, with the RBDs remaining in the “down” position (Figure 1A) ^6, 38^. On the other hand, a spontaneous large-scale conformational change occurs in the active CoV-1 spike protein simulation (Figure 1B), with the RBD moving from an active “up” position to a pseudo-inactive “down” conformation that is distinctly different from the inactive conformation in the cryo-EM structure ^38^. This spontaneous conformational transition appears to occur due to interactions between the NTD and RBD of the CoV-1 spike protein (Figure 1B). Unlike the active CoV-1 spike protein, the active CoV-2 spike protein does not undergo any large-scale conformational transitions and remains relatively stable within the 5-*µ*s simulations (Figure 1B). The RBD of the active CoV-2 spike protein remains in the “up” position (Figure 1B) ^6, 38^.

**Figure 1:**
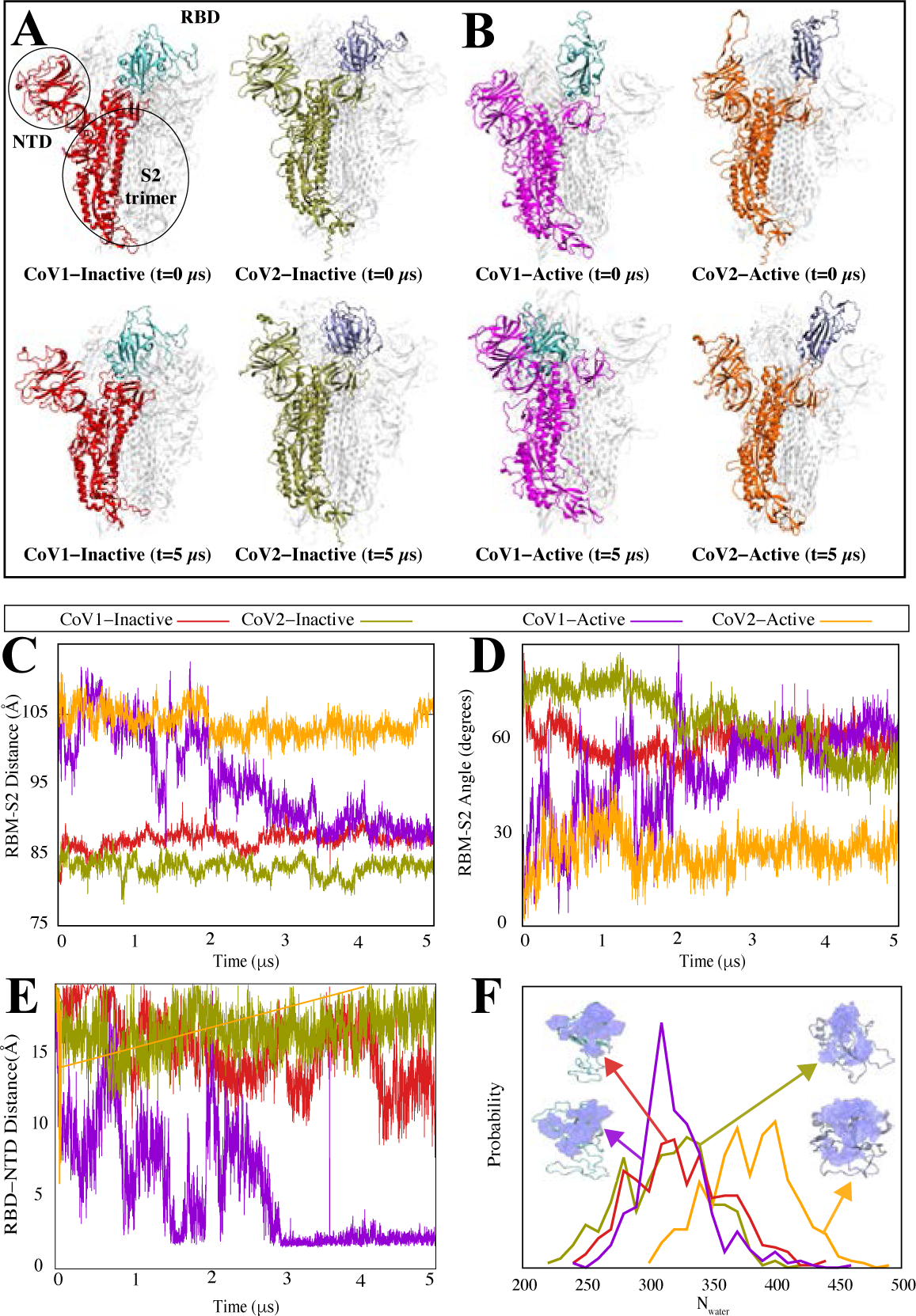
Unbiased simulations of the CoV-1 and CoV-2 spike proteins show a differential dynamic behavior. **(A-B)** The initial and final MD snapshots of CoV-1 and CoV-2 spike proteins starting from both inactive and active states. Protomer A in each protein is colored and protomers B and C are shown in white. The RBD of the colored protomer has a distinctive color from the rest of the protomer. Based on multiple repeats of these simulations, we have observed that the active form of the CoV-2 spike protein is consistently more stable than the active CoV-1 spike protein. The active CoV-1 spike protein transitions spontaneously to a pseudo-inactive conformation. **(C)** The center-of-mass distance between the S2 trimer of the spike protein and the RBM of protomer A shown as a function of time. **(D)** The angle between the S2 trimer of the spike protein and RBM of protomer A shown as a function of time. **(E)** Minimum distance between the NTD and RBD of protomer A as a function of time for CoV-1 and CoV-2 spike proteins in both active and inactive state simulations. **(F)** Probability density map of water within 5 Å of the RBM for the final 500 ns of simulation. In panels C-F, the same color code is used to represent CoV-1-inactive (blue), CoV-1-active (magenta), CoV-2-inactive (red) and CoV-2-active (orange).

To examine the reproducibility of the above observations, the active CoV-1 and active CoV-2 simulations were repeated two more times. Consistent with Set 1, the active CoV-2 simulations do not show any significant conformational change in Sets 2 and 3. The active CoV-1 simulations, on the other hand, undergo some significant conformational change in Set 2 and Set 3; although these conformational changes are not consistently observed in the three different repeats. The dramatic change from the “up” to “down” (or pseudo-inactive) conformation of the CoV-1 spike protein is only observed in Set 1; however, all three sets show some significant conformational changes that are not observed in any of the CoV-2 simulations. Root mean square deviation (RMSD) (Figure S1) and root mean square fluctuation (RMSF) (Figure S2) analyses demonstrate the relative stability of the active CoV-2 spike protein as compared to the active CoV-1 spike protein. A comparison of individual protomer RMSDs from all 3 replicas of the active CoV-1 and CoV-2 spike protein trajectories, clearly shows that the active CoV-1 spike protein is relatively less stable overall than the active CoV-2 spike protein (Figure S1). Similarly, RMSF analysis indicates that the RBD and NTD regions of the active CoV-1 spike protein fluctuate more than the corresponding regions of the active CoV-2 spike protein (Figure S2).

In order to quantify the spontaneous conformational transition that occurs in the active CoV-1 spike protein, we measured the center-of-mass distance between the receptor-binding motif (RBM) of protomer A and the S2 trimer of the spike protein (Figure 1C). The RBM-S2 distance remains stable for both inactive states at approximately 85 Å over 5 *µ*s. For both the CoV-1 and CoV-2 active states, the RBM-S2 distance is initially around 100 Å but decreases to approximately 85 Å for CoV-1 after 2 *µ*s (Figure 1C). This analysis clearly demonstrates that the final conformation adopted by the RBD of the active CoV-1 spike is similar to the inactive state RBD conformations of both CoV-1 and CoV-2, in terms of the RBM-S2 trimer distance (Figure 1C). On the other hand, the RBM-S2 trimer distance for the active CoV-2 spike protein remains relatively unchanged over 5 *µ*s (Figure 1C), consistent with the molecular images shown in Figs. 1A-B. Similarly, the angle between the RBM of protomer A and the S2 trimer remains relatively unchanged for the CoV-2 active state, while the CoV-1 active simulation shows a behavior during the last 3 *µ*s that is similar to that of the inactive states of CoV-1 and CoV-2 (Figure 1D).

We also calculated the minimum distance between the RBD and NTD of protomer A for each system (Figure 1E), which quantifies the motion and position of the RBD relative to the NTD. While the RBM-S2 distance and angle calculations indicate that the behavior of the CoV-1 active state eventually resembles that of both inactive systems (Figure 1C-D), the NTD-RBD distance calculation showcases the unique behavior of the active CoV-1 spike protein. The NTD-RBD distance of the active protomer in CoV-1 fluctuates considerably over the first 2 *µ*s of the trajectory, after which it decreases sharply to settle down between 1-2 Å (Figure 1E). This clearly demonstrates that the RBD of the active CoV-1 spike protein is in close proximity to the NTD as observed during a visual inspection of the trajectories (Figure 1B). This is not observed for the active CoV-2 spike protein or either of the inactive spike proteins (Figure 1A-B, 1E), thus indicating that the pseudo-inactive conformation adopted by the active CoV-1 spike protein is unique. Additionally, a probability density map was generated for water molecules within 5 Å of the RBM during the last 500 ns of simulation (Figure 1F). The water molecule count for the CoV-1 active state is lower than that of the CoV-2 active state and is comparable to the counts for the CoV-1/2 inactive states, further confirming that the active CoV-1 spike protein undergoes a large-scale conformational transition (Figure 1F).

### Principal component analysis and dynamic network analysis provide evidence of the conformational stability of the active CoV-2 spike protein

We performed principal component analysis (PCA) to validate our claim that the active form of the CoV-2 spike protein is more stable than the active CoV-1 spike protein and to provide insight into the mechanistic aspects of the spike protein activation-inactivation process. When the individual protomer trajectories (see Methods section) from the CoV-1/CoV-2 active (Set 1) and inactive simulations are projected onto the space of their first two principal components (PC1 and PC2), it clearly demonstrates that the CoV-1 active protomer A samples a much larger region in the PC1 space than CoV-2 active protomer A (Figure 2A, 2C). This is further evidence of the relative stability of the active CoV-2 spike protein in comparison to the active CoV-1 spike protein.

**Figure 2:**
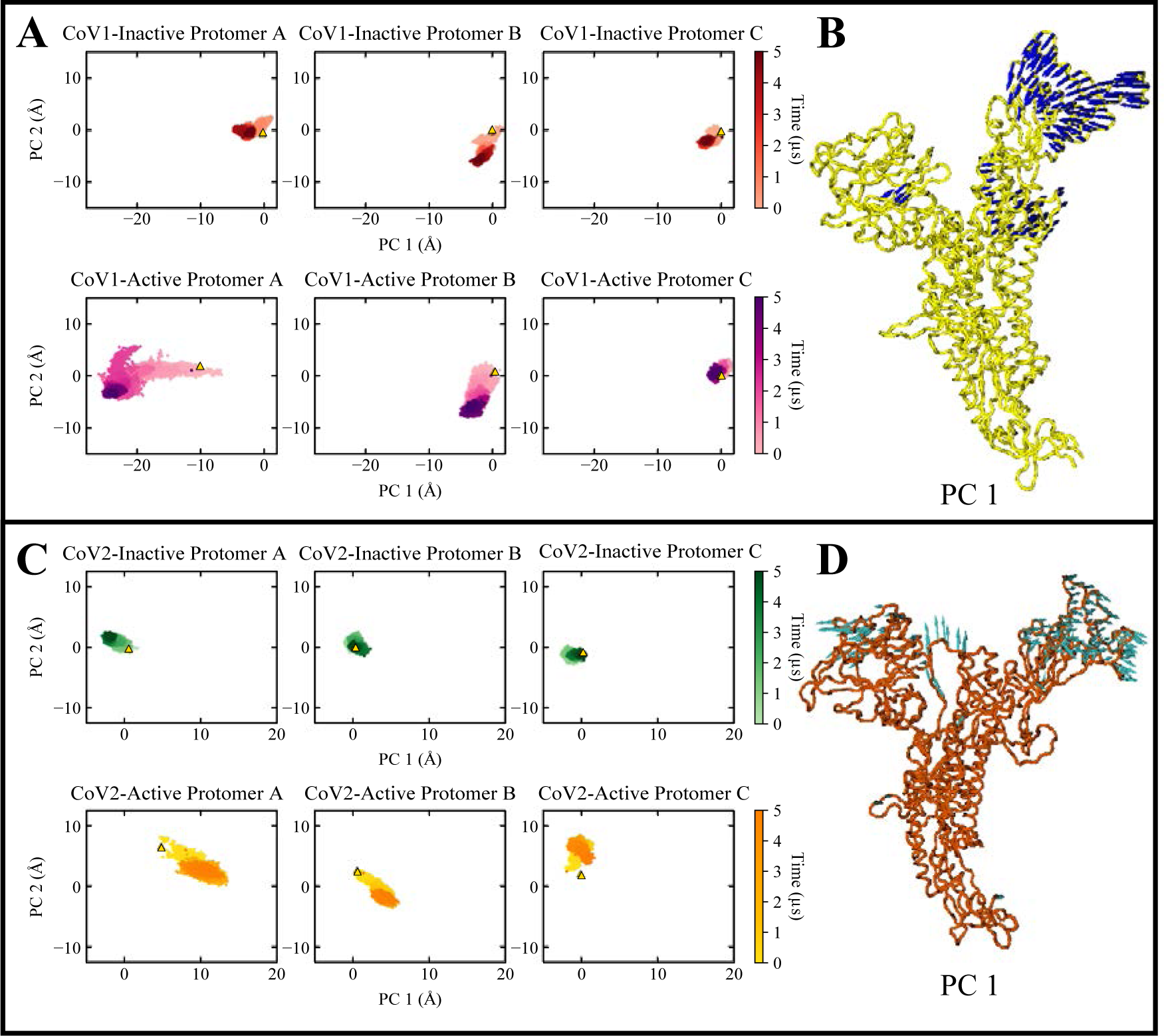
Principal component analysis demonstrates that the active CoV-2 spike protein is more stable than the active Cov-1 spike protein. (A) Scatter plot of PC1 and PC2 for each protomer in the active and inactive CoV-1 simulations. Protomers from inactive state simulations are colored red while protomers from active state simulations are colored magenta. Lighter/darker colors represent earlier/later stages in the simulation. (B) Visual representation of PC1 with the blue arrows at each C-*α* atom indicating direction and magnitude of variance. The RBD of the CoV-1 spike protein shows pronounced motions in the direction of the NTD. (C) Scatter plot of PC1 and PC2 for each protomer in the inactive and active CoV-2 simulations. Protomers from inactive state simulations are colored green while protomers from active state simulations are colored yellow. The active CoV-2 spike protein is relatively stable and samples significantly fewer conformations in the PC1 space in comparison to the active Cov-1 spike protein. (D) Visual representation of PC1 with the cyan arrows at each C-*α* atom indicating direction and magnitude of variance. The NTD and RBD of the CoV-2 spike protein show slight movement away from each other.

A visual representation of PC1 for all protomers from the CoV-1 spike protein simulations shows that the RBD undergoes the most pronounced motions directed inward towards the NTD (Figure 2B). On the other hand, a visual representation of PC1 for the CoV-2 spike protein shows that the RBD and NTD tend to move away from each other slightly and that the fluctuations are significantly smaller than in the CoV-1 spike protein (Figure 2D). The most pronounced collective motion in each system (PC1) describes the distinct motions associated with the RBD, that play key roles in the inactivation of the active CoV-1 spike protein and maintenance of the active conformation of the CoV-2 spike protein (Figure 1). This highlights the differential dynamic behavior of the active CoV-1 spike protein.

PC2 describes the relative motions of the NTD and RBD, showing that the NTD motion is more pronounced in CoV-1 (Figure S3). The motions associated with PC2 are roughly the opposite of those associated with PC1 in terms of direction. PC2 also shows that the CoV-1 spike protein has more regions outside the NTD and RBD that show high variance (Figure S3). Similar trends are observed in Sets 2 and 3 of the active state simulations (Figure S4). While different protomers are involved, the active CoV-1 spike protein still undergoes more pronounced motions in both PC1 and PC2 compared to the active CoV-2 spike protein (Figure S4). These observations are in agreement with our claim that the active CoV-2 spike protein is relatively stable and that the active CoV-1 spike protein transitions spontaneously to a pseudo-inactive conformation.

The inferences drawn from PCA are also supported by dynamic network analysis (DNA). Differential behavior of the active CoV-1 and CoV-2 spike proteins manifests in the correlation of motions between the various domains in individual protomers. In Figure 3A, correlation heat maps of active CoV-1 protomer A (Set 1) and inactive CoV-1 protomer C are presented, along with the difference between the active state and the reference structure (inactive protomer C). The heat map for active Cov-1 protomer A shows regions of high correlation and anticorrelation between several domains of the protomer. The NTD correlates strongly with itself while anticorrelating with the RBD and parts of the S2 region. The reference protomer, inactive CoV-1 protomer C, shows a general reduction in correlation across all regions (Figure 3A). The NTD does correlate with itself, but not as strongly as in the active CoV-1 protomer A. Similarly, the NTD-RBD anticorrelations were reduced. The Δ matrix of differences between active CoV-1 protomer A and inactive protomer C identified the regions where the correlations were most different. Correlations between S1-C and the NTD/RBD changed significantly, as did correlations between the RBD and S2 region (Figure 3A).

**Figure 3:**
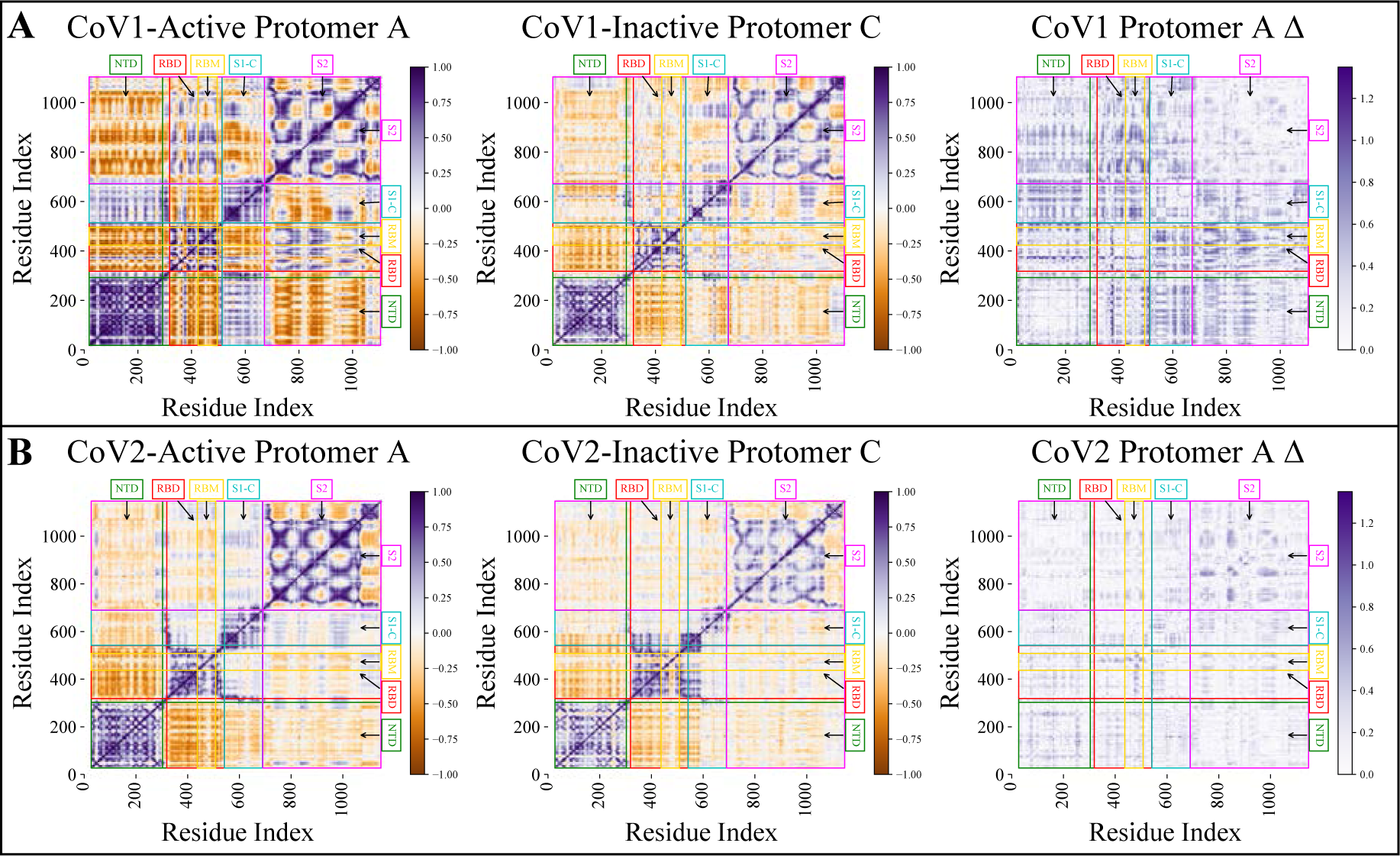
Dynamic network analysis shows that intra-protomer correlations and anticorrelations are relatively strong in the active CoV-1 spike protein simulations. **(A)** DNA heat maps showing the correlation of motions for the active CoV-1 protomer A, inactive protomer C (reference), and the difference matrix. **(B)** DNA heat maps showing the correlation of motions for the active CoV-2 protomer A, inactive protomer C (reference), and the difference matrix. Correlations are shown in purple and anti-correlations are shown in orange, with the darker colors indicating greater correlation/anti-correlation. Colored labels for the NTD (green), RBD (red), RBM (yellow), S1-C (cyan), and S2 (magenta) regions are positioned over the appropriate residues. The delta matrix identifies differences in protomer correlation between the active and reference inactive protomer. A theoretical maximum for Δ is 2, but the observed maximum was less than 1.3. Differences in correlation are shown as a purple gradient with darker purple indicating larger difference.

The correlations and anti-correlations observed for active CoV-2 protomer A (Set 1) were not as strong as those observed for active CoV-1 protomer A (Figure 3B). Similar to CoV-1, anti-correlation occurs between the NTD and RBD but is not as pronounced. Very low correlation was observed between the NTD and S1-C/S2 regions, also differentiating CoV-2 from CoV-1. The active CoV-2 protomer A is closer to the stable inactive CoV-2 protomer C, as shown in the Δ matrix (Figure 3B). DNA correlation heat maps for all protomers in Set 1 of the CoV-1/CoV-2 active state simulations may be found in the supporting information (Figure S5-S6). Similar trends were observed in Set 2 and Set 3 of the CoV-1 and CoV-2 active state simulations, shown in Figure S7 and S8 respectively. These observations thus provide further evidence of the relative stability of the active CoV-2 spike protein.

The concerted movements of each protomer relative to the rest of the trimer also highlight the differences between the active CoV-1 and CoV-2 spike proteins. Heat maps showing correlations between NTD regions of different protomers are presented in Figure 4A. Stronger correlations and anticorrelations occurred in Sets 2 and 3 of the active CoV-1 simulations (Figure 4A). Set 2 showed moderately strong anticorrelations between NTDs A-C and NTDs B-C. Stronger anti-correlations between NTDs A-B and NTDs B-C occurred in Set 3, with moderate correlations between NTDs A-C. The active CoV-2 simulations showed similar correlations across all three simulation sets, with slightly increased values in Set 3 (Figure 4A). These observations are consistent with a more stable conformation for the active CoV-2 spike protein.

**Figure 4:**
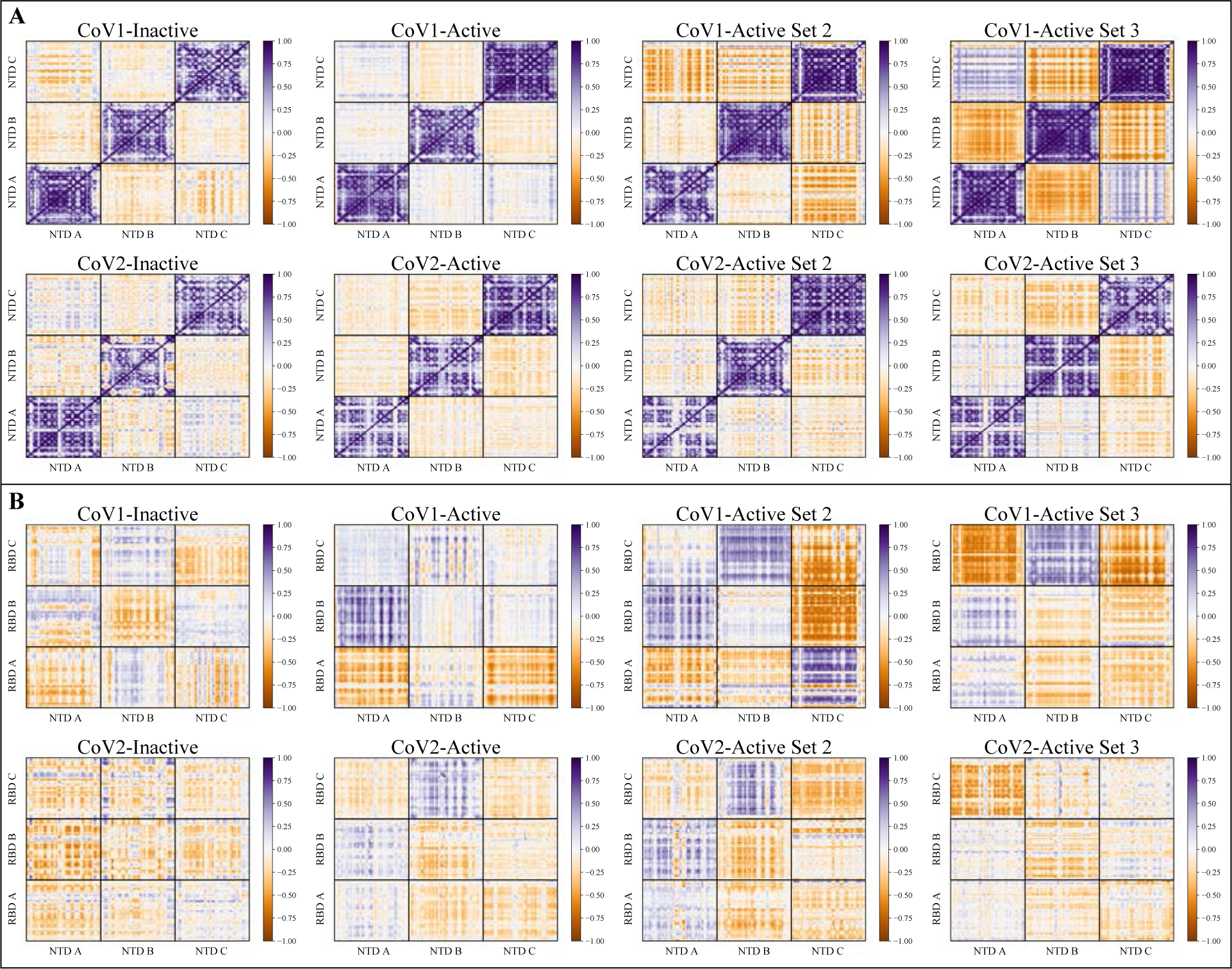
Dynamic network analysis shows that inter-protomer correlations and anticorrelations are relatively strong in the active CoV-1 spike protein simulations. **(A)** DNA heat maps showing the correlation of motion between the NTD regions of different protomers. **(B)** DNA heat maps showing the correlation of motion between the NTD and RBD regions of different protomers. Correlations are shown in purple and anti-correlations are shown in orange, with the darker colors indicating greater correlation/anti-correlation.

Figure 4B shows a similar trend with correlations between the NTD and RBD regions of different protomers. Sets 2 and 3 of the active CoV-1 spike protein trajectories showed stronger correlations between the NTD and RBD regions than the corresponding CoV-2 trajectories (Figure 4B). In particular, RBD C of Sets 2 and 3 had strong correlations or anticorrelations with the NTDs of all protomers (Figure 4B). The CoV-2 simulations displayed lower correlations for all the NTD-RBD combinations, with similar results for both active state and inactive state trajectories (Figure 4B). This recapitulates our other observations of greater conformational stability of the active CoV-2 spike protein relative to the active CoV-1 spike protein (Figure 1-3).

### Inter-domain electrostatic interactions between the NTD and RBD drive the inactivation of the active CoV-1 spike protein

As described previously, the active CoV-1 spike protein undergoes a spontaneous large-scale conformational transition and essentially becomes inactivated (Figure 1A-F). The RBD of the active protomer A moves towards and gets very close to the NTD, which does not occur in the CoV-2 active and CoV-1/2 inactive trajectories (Figure 5A-B). The driving force behind this conformational transition is a set of salt-bridge interactions that are unique to the active CoV-1 spike protein. Residues D23 and D24 in the NTD interact with K365 in the RBD, forming stable salt bridges in the active CoV-1 spike protein but not in the inactive state (Figure 5C-D). These fairly stable salt-bridges form around the 1 *µ*s mark (Figure 5C-D), prior to the final movement of the RBD towards the NTD which happens after 2 *µ*s (Figure 5A). Residues D23 and D24 are not conserved in the SARS-CoV-2 spike protein.

**Figure 5:**
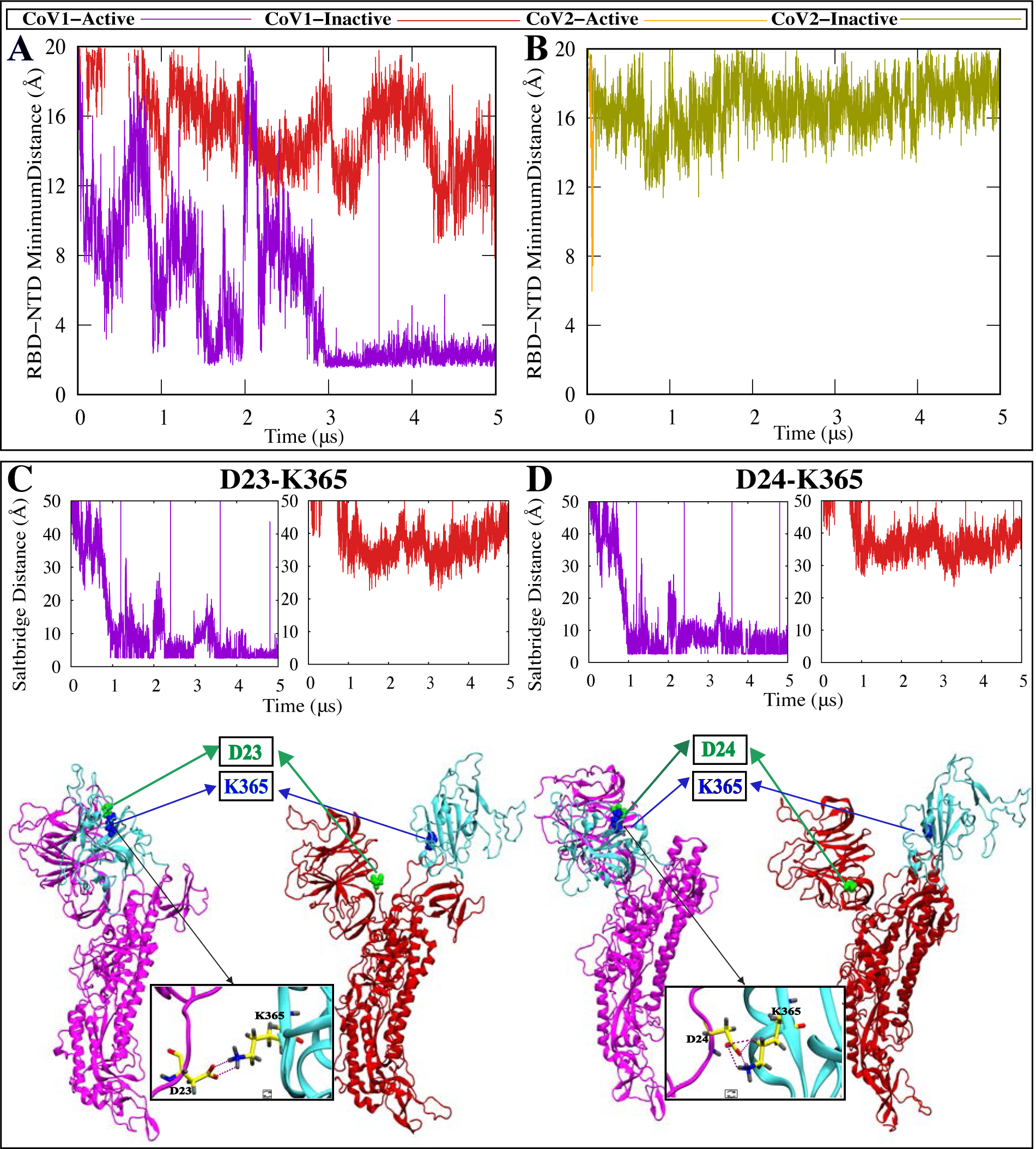
Unique salt-bridge interactions between the RBD and NTD of the active CoV-1 spike protomer facilitate the transition to a pseudo-inactive conformation. **(A-B)** Minimum contact distance between the RBD and NTD of protomer A, in the CoV-1 and Cov-2 spike proteins respectively. The RBD of active CoV-1 protomer A moves towards and interacts with the NTD. **(C-D)** In the active CoV-1 protomer A, D23 and D24 (green) of the NTD form a salt-bridge with K365 (blue) of the RBD. This salt-bridge interaction is absent in the inactive protomers. D23 and D24 are not conserved in the CoV-2 spike protein. Time series of D23/24-K365 salt-bridge distances and visual representations of salt-bridge formation.

Differential behavior is also observed for two sets of residues that are conserved in both CoV-1 and CoV-2 spike proteins (Figure S9). R328 and D578 form a stable salt bridge in both active and inactive CoV-2 spike proteins while R315 and D564 do not form a salt-bridge in the CoV-1 spike proteins (Figure S9A). Similarly, R273 and D290 form a stable salt bridge in the both active and inactive CoV-2 spike proteins while K258 and D277 do not form a salt-bridge in the CoV-1 spike proteins (Figure S9B). Additionally, a conserved pair of residues form an intra-RBD hydrogen bond in the active/inactive CoV-2 spike protein (Y396-E516) and the inactive CoV-1 spike protein (Y383-E502), but not in the active CoV-1 spike protein (Y383-E502) (Figure S10A-B). These electrostatic interactions thus potentially contribute to the relative stability of the active SARS-CoV-2 spike protein. These observations are in agreement with the results obtained through visual inspection of the trajectories as well as the other analyses described previously (Figure 1-4).

The propensity of the active CoV-1 spike protein to deviate from its ”RBD-up” conformation is in marked contrast to the stability displayed by the active CoV-2 spike protein in our unbiased microsecond-level simulations. The active SARS-CoV-1 spike protein consistently exhibits a differential dynamic behavior that could potentially explain why SARS-CoV-2 is more transmissible than SARS-Cov-1 ^8, 9^.

### SMD simulations indicate that the SARS-Cov-2 spike protein has larger energy barriers to large-scale conformational changes than the CoV-1 spike protein

SMD simulations were performed to quantify and characterize the energetics of the activation-inactivation process for the SARS CoV-1 and CoV-2 spike proteins. To induce the activation or inactivation of individual protomers, we used collective variables based on the C*α* RMSD of each protomer. 10 sets of 100 ns SMD simulations were performed for each system. The conformational transition of an inactive receptor-binding domain to the active “up” position was accompanied by a decrease in the RBM-S2 angle and an increase in the RBM-S2 distance (Figure 6A-B). Similarly, the inactivation of an active protomer was characterized by an increase in the RBM-S2 angle and a decrease in the RBM-S2 distance (Figure 6A-B).

**Figure 6:**
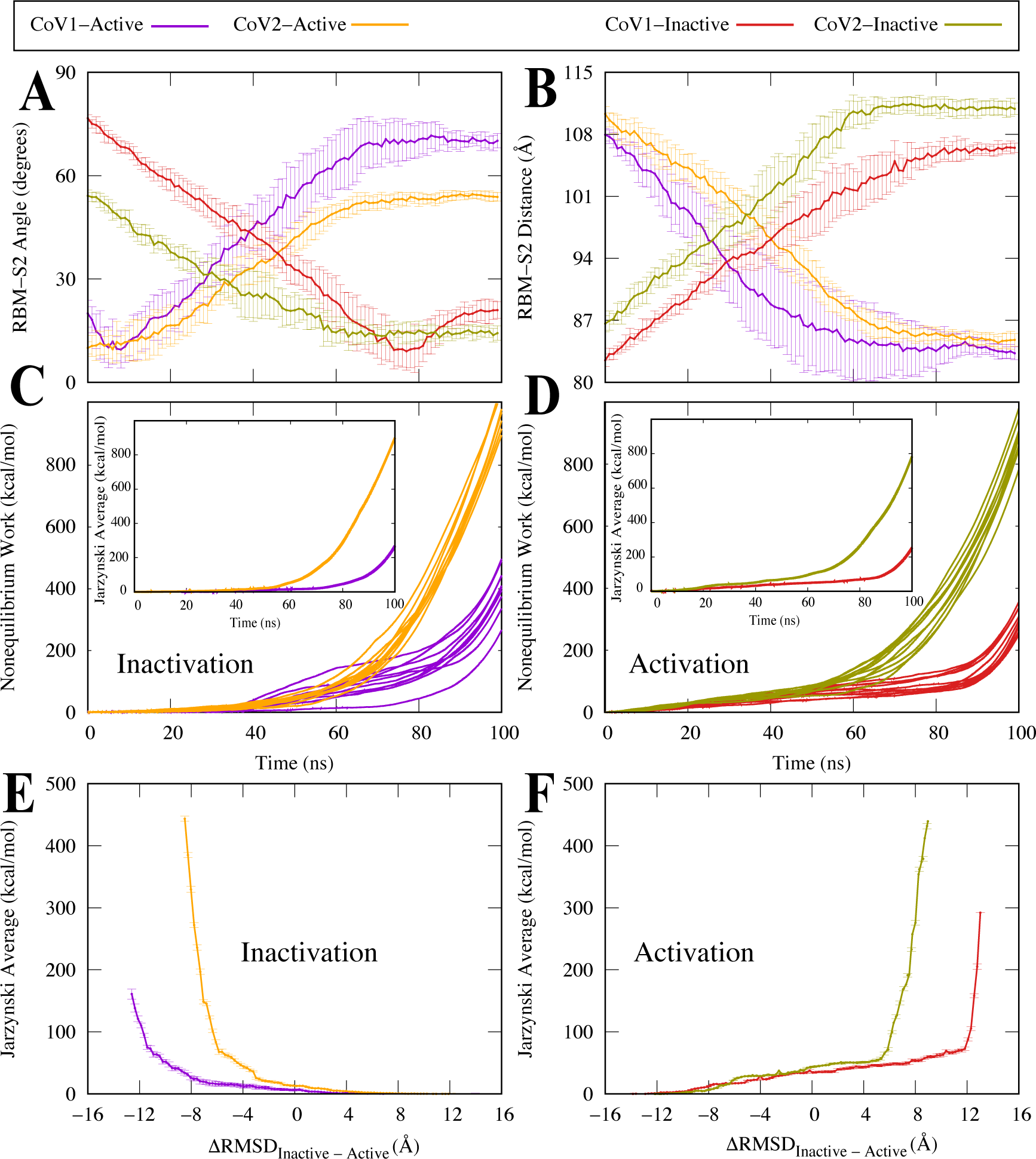
SMD simulations show that the CoV-2 spike protein has higher energy barriers between active and inactive states than the CoV-1 spike protein. **(A)** RBM-S2 Angle between the beta sheet region of the RBM and the alpha helical region of S2, shown as a function of time. Protomer activation is characterized by a decrease in the RBM-S2 angle. **(B)** RBM-S2 COM Distance between the beta sheet region of the RBM and the alpha helical region of S2, shown as shown as a function of time. Protomer activation is characterized by an increase in the RBM-S2 distance.**(C,D)** Accumulated non-equilibrium work performed by a single protomer as a function of time. **(E,F)** Jarzynski average taken with respect to the Δ*RMSD* = *RMSD_Inactive_* - *RMSD_Active_*.

Without performing strict free-energy calculations, we have used non-equilibrium work measurements to compare the thermodynamics and kinetics of the CoV-1 and CoV-2 spike protein activation-inactivation process in a semi-quantitative manner. We have previously used similar methods to investigate conformational transitions of other biomolecular systems ^50–53^. The accumulated non-equilibrium work measured during the inactivation of an active CoV-2 protomer or the activation of an inactive CoV-2 protomer, is significantly larger than the work measured during the inactivation or activation of a CoV-1 protomer (Figure 6C-D). Similarly, the change in the associated Jarzynski average is also much higher for the CoV-2 protomers (Figure 6C-D). These results strongly suggest that the CoV-2 spike protein has slower kinetics in both directions and that the conformational change associated with activation or inactivation proceeds more slowly than in the Cov-1 spike protein. This is in very good agreement with our observations on the relative conformational stability of the active CoV-2 spike protein from the unbiased simulations.

Additionally, the change in Jarzynski average with respect to the difference in RMSD between inactive and active states, is relatively higher for the CoV-2 spike protein (Figure 6E-F). The Jarzynksi average associated with inactivation of the active CoV-1 spike protein is much lower than the Jarzynski average associated with the other conformational transitions (Figure 6E). This indicates that the inactivation process is much easier for the CoV-1 spike protein in comparison to the CoV-2 spike protein and explains why we were able to observe a spontaneous conformational transition in the unbiased active CoV-1 trajectory within 5 *µ*s. On the other hand, there are considerable energy barriers to the activation process for both CoV-1 and Cov-2 spike proteins (Figure 6F), explaining the fact that the inactive forms of both spike proteins remain extremely stable during our unbiased simulations. These results from our biased SMD simulations are thus very consistent with the results from our unbiased equilibrium simulations, highlighting the stability of the active CoV-2 spike protein and the relative conformational plasticity of the active CoV-1 spike protein.

Our SMD simulations thus conclusively show that it is relatively difficult for the CoV-2 spike protein to undergo a large-scale conformational transition between active and inactive states, when compared to the CoV-1 spike protein. These SMD simulations involved single protomers and were also repeated with all 3 protomers (Figure S11). We observed similar behavioral trends in the multi-protomer SMD simulations (Figure S11). Our results indicate that the energy barriers to these conformational changes are larger in the CoV-2 spike protein and that the activation-inactivation mechanism might be more energetically favorable in the CoV-1 spike protein. This provides a rationale for the spontaneous conformational transition observed in the active CoV-1 spike protein equilibrium simulation and the absence of similar conformational changes in the corresponding CoV-2 spike protein simulation (Figure 1).

## Discussion

Using microsecond-timescale unbiased and biased simulations, we have demonstrated that the active SARS-CoV-2 and SARS-CoV-1 spike proteins exhibit differential dynamic behavior. The active CoV-2 spike protein remains relatively stable over 5 *µ*s, whereas the active CoV-1 spike protein spontaneously adopts a pseudo-inactive conformation that is distinct from the well-characterized inactive “RBD-down” conformation ^38^. Our discovery of a pseudo-inactive state of the CoV-1 spike protein essentially agrees with the results of an experimental smFRET study that describes alternative inactive states of the CoV-2 spike protein ^48^. While this pseudo-inactive conformation is not observed in our CoV-2 spike protein simulations, it is certainly plausible that the CoV-2 spike protein samples alternative conformational states during the spike protein activation process that is dependent on the experimental/physiological conditions.

An interesting feature of the pseudo-inactive conformation of the CoV-1 spike protein is the interaction between the RBD and NTD, which is not observed in the previously known inactive conformation ^38^. As shown by PCA analysis, the RBD of the active CoV-1 spike protein moves inward towards the NTD. This pronounced motion of the RBD enables the formation of unique salt-bridge interactions between the NTD and RBD, which drive the conformational transition. We have also identified stabilizing salt-bridge and hydrogen-bond interactions between conserved residue pairs, that form in the CoV-2 spike protein but not in the CoV-1 spike protein.

In general, our PCA analysis indicates that the active CoV-1 spike protein shows more pronounced motions over the course of the simulations. This is corroborated by dynamic network analysis, which illustrates that the active CoV-2 spike protein has markedly weaker intra-protomer and inter-protomer correlations and anticorrelations than the active CoV-1 spike protein. These observations are consistent with the relative conformational stability of the active CoV-2 spike protein. An investigation of the energetics of the activation-inactivation process using SMD simulations revealed that relative to CoV-1, it is difficult for the CoV-2 spike protein to undergo a major conformational transition from the active state to the inactive state or vice-versa. Non-equilibrium work measurements indicate that large-scale conformational transitions occur relatively slowly in the CoV-2 spike protein, which complements our observations on the relative conformational stability of the active CoV-2 spike protein from the equilibrium simulations. We also found that the energy barriers involved in the inactivation of the CoV-1 spike protein are quite low, thus explaining the spontaneous conformational transition observed in the active CoV-1 equilibrium trajectory. The results from our equilibrium and non-equilibrium simulations are thus very consistent and provide extensive insights into the long-term dynamics of the CoV-1 and CoV-2 spike proteins. A recent computational study has shown that the RBD of the CoV-2 spike protein has greater mechanical stability than the RBD of the CoV-1 spike protein ^54^, which agrees with our observations on the conformational stability of the active CoV-2 spike protein.

Recently, Anand et al. proposed that the SARS-CoV-1 spike protein needs to overcome comparatively higher energy barriers to adopt the active conformation ^55^, which is the opposite of what we observe in our simulations. However, their proposed energy landscape is based on inferences drawn from cryo-EM data and does not take the inherently dynamic nature of the proteins into account ^55^. Unlike X-ray crystallography and cryo-EM, MD simulations are able to provide detailed information on the dynamic behavior of proteins and other biomolecules ^46, 47^. However, each computational or experimental technique has its own assumptions and limitations. Here, for instance, we chose to work with the non-glycosylated spike proteins of CoV-1 and 2 to avoid complications in comparisons. A recent study has shown that glycosylation of the spike proteins might play an important role in the conformational dynamics of the RBD ^56^. At this stage, we have not simulated the glycosylated spike proteins due to the difficulty of modeling the correct glycan chains. It would be quite difficult to determine whether conformational changes occur as a result of the intrinsic protein dynamics or the differential glycosylation patterns of the CoV-1 and CoV-2 spike proteins imposed by our modeling. However, we use the non-glycosylated form of the spike protein for both CoV-1 and 2, which makes the comparison justifiable.

Our simulations thus provide valuable insight into the dynamic behavior of the CoV-1 and CoV-2 spike proteins. An improved understanding of the conformational changes leading to activation or inactivation of the spike proteins, is critical to the effective development of novel therapeutics and vaccines using a structure-based drug design framework. As discussed earlier, investigation of the ”effective binding” process involving both receptor interaction and spike protein activation will provide deeper insights into the enhanced infectivity of SARS-Cov-2. Several studies have investigated RBD-ACE2 binding for both SARS CoV-1 ^57–63^ and SARS-CoV-2 ^6, 17, 27, 36, 37^, while ignoring the conformational dynamics of spike protein activation and inactivation. For the first time, our microsecond-level unbiased and biased simulations provide insights into the differential activation and inactivation processes of the SARS-CoV-1 and CoV-2 spike proteins. Therefore, we propose that the ”effective binding” process is different in the CoV-1 and CoV-2 spike proteins, not only because of the variability of the RBD but also due to the contribution of other regions like the NTD (as seen in the spontaneous conformational change involving the active CoV-1 spike protein). Our simulations strongly suggest that the differential conformational dynamics associated with inactivation and activation might contribute to the increased transmissibility of SARS-CoV-2. However, additional experimental and computational studies are needed to fully investigate this possibility.

## Methods

### All-atom equilibrium MD simulations

We have used all-atom equilibrium MD simulations to characterize the conformational dynamics of the spike protein from SARS-CoV-2 and SARS-CoV-1. Our simulations were based on cryo-EM structures of the SARS-CoV-2 spike protein in the active (PDB entry:6VYB) ^6^ and inactive (PDB entry:6VXX) ^6^ states and the SARS-CoV-1 spike protein in the active (PDB entry:5X5B) ^38^ and inactive (PDB entry:5X58) ^38^ states. Missing residues and loop regions for all 4 models were generated using Modeller ^64^. A Monte Carlo algorithm was used to iteratively minimize the energy of the system ^64^. 10,000 Monte Carlo iterations were used to generate the initial models for the equilibrium simulations. CHARMM-GUI ^65, 66^ was then used to build the simulation systems. Engineered residues (P986, P987) ^41, 67^ in the SARS-CoV-2 spike protein were mutated back to the wildtype residues (K986, V987). The protein was solvated in a box of TIP3P waters, and 0.15 M NaCl (in addition to the counterions used to neutralize the protein) using CHARMM-GUI ^65, 66^. The box size for the CoV-2 active model was 198 x 198 x 198 Å^3^ with 730937 atoms. The box size for the CoV-2 inactive model was 183 x 183 x 183 Å^3^ with 577927 atoms. The box size for the CoV-1 active model was 193 x 193 x 193 Å^3^ with 680615 atoms. The box size for the CoV-1 inactive model was 169 x 169 x 169 Å^3^ with 454608 atoms.

All simulations were performed using the NAMD 2.13 ^68^ simulation package with the CHARMM36 all-atom additive force field ^69^. The input files for energy minimization and production were generated using CHARMM-GUI ^65, 66^. Initially, we energy-minimized each system for 10,000 steps using the conjugate gradient algorithm ^70^. Then, we relaxed the systems using restrained MD simulations in a stepwise manner using the standard CHARMM-GUI protocol ^65, 66^ (”relaxation step”). In the next step, backbone and sidechain restraints were used for 10 ns with a force constant of 50 kcal/mol.Å ^2^ (”restraining step”). The systems were then equilibrated with no bias for another 10 ns (”equilibration step”). The initial relaxation was performed in an NVT ensemble while the rest of the simulations were performed in an NPT ensemble. Simulations were carried out using a 2-fs time step at 310 K using a Langevin integrator with a damping coefficient of *γ* = 0.5 ps*^−^*1. The pressure was maintained at 1 atm using the Nose-Hoover Langevin piston method ^70, 71^. The smoothed cutoff distance for non-bonded interactions was set at 10 to 12 Å and long-range electrostatic interactions were computed with the particle mesh Ewald (PME) method ^72^.

These initial simulations were executed on TACC Longhorn. The production run for each model was then extended to 5 *µ*s on Anton2 ^73^, with a timestep of 2.5 fs. Conformations were collected every 240 picoseconds. Initial processing of the Anton2 simulation trajectories was carried out on Kollman ^73^. Two additional 5 *µ*s simulations were performed for both the CoV-2 and CoV-1 active models on Anton2 (referred to as Set 2 and Set 3 in the manuscript). The initial production runs for these models on TACC Longhorn were extended twice by 0.5 ns in order to generate the starting conformations for the repeat simulations. 40 *µ*s of simulation data was generated in aggregate – 15 *µ*s each for the active Cov-1/Cov-2 spike proteins and 5 *µ*s each for the inactive spike proteins.

### RBM-S2 distance and angle calculations

To quantify the RBM-S2 distance, we defined centers of mass based on residues that form a beta-sheet in the RBM region of each RBD (CoV-1: RBM residues 439 to 441, 479 to 481; CoV-2: RBM residues 452 to 454, 492 to 494) and residues that encompass the S2 trimer (CoV-1: S2 residues 672 to 1104; CoV-2: S2 residues 690 to 1147). We then measured the vector distance between the two centers of mass and used the vector magnitude to quantify the overall distance.

For the RBM-S2 angle, we chose residues at the top and bottom of the straightest region of the S2 Trimer (alpha-helical regions in CoV-1: residues 970 and 1016; CoV-2: residues 914 and 987). Similarly, we also chose residues from the beta-sheet region of the RBM and one at the bottom of the RBD (CoV-1: residues 348 and 478; CoV-2: residues 391 and 493). We then defined a vector direction using the vector subtraction of the two chosen residues in the S2 region and the residues of the RBD region, which were defined as *v*_1_ and *v*_2_. The vector angle between the RBD and S2 was then calculated with the following equation: arccos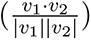. The computed angle was subtracted from 180*^◦^*. An angle above *≈*60^◦^ would indicate an RBD in the inactive conformation with respect to S2, and 0-40*^◦^* would indicate an RBD in the active conformation.

### NTD-RBD distance calculations and water count

To characterize conformational changes in the active and inactive states of both CoV-1 and CoV-2 spike proteins, we calculated the minimum distance between every residue of the receptor-binding domain (RBD) and the N-terminal domain (NTD). We measured the distance between each residue pair in these regions (maximum distance cut off was 20 Å) as a function of time. The domains were defined as follows: CoV-2 RBD (residues 330 to 515); CoV-2 NTD (residues 60 to 270); CoV-1 RBD (residues 330 to 550); CoV-1 NTD (residues 35 to 255).

The amount of solvent around the receptor-binding motif (RBM) was quantified using a VMD^74^ script. We calculated the number of water molecules within 5 Å of the RBM for every frame of the last 500 ns of each trajectory and also plotted probability density maps for each water count.

### Principal Component Analysis (PCA)

PCA performed with ProDy ^75^ was used to quantify the persistent conformational changes and relative motions of the active and inactive states. Only the position of the C-*α* atoms of the spike protein was considered when building the covariance matrix of atomic positions, in order to focus on the large conformational changes and ignore side chain fluctuations. Each trajectory was aligned with the positions from the cryo-EM structure before analysis to remove translational motion of the protein from the variance calculations.

The CoV-1/CoV-2 active state (Set 1) and CoV-1/CoV-2 inactive state trajectories were stripped down to trajectories of the individual protomers from each simulation. The individual protomers were then analyzed together to compare and quantify the relative motions of the active and inactive states. Through eigenvalue decomposition, the top twenty principal components (PCs) were calculated for each protomer. The top two PCs for each protomer have been plotted to identify the major motions of the protein.

### Dynamic Network Analysis (DNA)

DNA of the correlated motions of the protein provided further quantitative information on the concerted motions of the C-*α* atoms of the protein. MD-TASK ^76^, a software suite of MD analysis tools, was used to calculate the correlation coefficient for the motion of each C-*α* atom relative to the other C-*α* atoms. A correlation matrix *M* was generated for each of the three protomers in all the simulated trajectories. Additionally, a correlation matrix for the entire trimer was calculated for each simulation to explore correlations between structures of different protomers. A step size of four frames was used during the correlation calculations to reduce the processing times, given the large number of residues.

To quantify the differences in correlation between a protomer and some reference, a difference matrix, Δ was calculated,

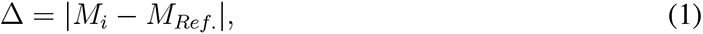

where *M_i_* is the correlation matrix of interest, and *M_Ref_* is the correlation matrix of a reference conformation. In this work, the difference between a protomer in an active conformation and an inactive conformation was of interest. For this reason, the protomers in the active simulations were compared with Protomer C in the inactive simulation, which displayed relatively little motion.

### Electrostatic interaction analysis

To identify interactions that contribute to the stability of the Cov-2 spike protein or play key roles in the CoV-1 active conformational transition, we performed salt-bridge and hydrogen-bond analysis for all SARS-CoV-2 and SARS-CoV-1 systems. Salt bridges were identified using the VMD Timeline plugin ^74^ at a cutoff distance of 4.0 Å . The salt-bridge cutoff distance is defined as the distance between the oxygen atom of the participating acidic residue and the nitrogen atom of the basic residue. The VMD HBond plugin ^74^ was used for hydrogen bond analysis. The donor-acceptor distance and angle cutoffs used were 3.5 Å and 30 degrees respectively. We report salt-bridge and hydrogen-bond interactions that illustrate the differential behavior of the SARS-CoV-2 and CoV-1 spike proteins.

### Steered Molecular Dynamics (SMD) Simulations

To induce activation/inactivation of a protomer initially in the inactive/active conformation, we defined collective variables based on the the C*α* RMSD of each protomer in the CoV-1 and CoV-2 systems. Reference coordinates were taken from the corresponding active/inactive structure for both CoV-1 and CoV-2 protomers. The atoms chosen were based on the total number of resolved and modeled residues in the CoV-2 structures. Structural analysis of CoV-1 and CoV-2 was employed to ensure that equivalent C*α* atoms were steered in all simulation sets. 1037 atoms were steered for any given protomer and the following residue range was used: 27 to 239, 244 to 315, 322 to 662, 673 to 809, and 831 to 1104. These atoms span the entire protomer, starting from the NTD and ending approximately at the C-terminus of the S2 region. A force constant of 250 *kcal/mol/*Å^2^ was used for SMD simulations involving a single protomer and a force constant of 750 *kcal/mol/*Å^2^ was used for SMD simulations involving all three protomers.

The systems used for each simulation were taken from the outcome of the ”equilibration step” (see Initial Preparation in Methods) as explained in the equilibrium simulation methods. Utilizing the multi-copy capabilities of NAMD, we performed 10 sets of 100 ns RMSD steering for each system - 8*µ*s of simulation time in aggregate.

For all SMD time series analyses, each data point was averaged for the 10 sets and standard deviation was calculated. Each analysis was plotted with 100 points and error bars were derived from the standard deviation. The RBM-S2 distance and angle calculations were performed as described previously.

Using the Jarzynski relation^77^ we calculate the Jarzynski average at time *t* during the activation or inactivation process as 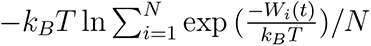, where *k_B_* and *T* are the Boltzmann constant and the temperature, respectively and *W_i_*(*t*) is the work accumulated from the beginning of the SMD simulation *i* up to time *t*. The above average would converge to the free energy for large number of trajectories (*N → ∞*). For *N* = 10, the above average simply provides a semi-quantitative measure for relative energetic comparisons ^50–53^.

All RMSD calculations are represented as the ΔRMSD. The ΔRMSD represents the RMSD with respect to the inactive conformation minus the RMSD with respect to the active conformation for all systems. We binned the ΔRMSD space and used all snapshots of each SMD trajectory to collect all work values associated with each bin. 100 bins were used. We then calculated the Jarzynski average for each bin. Standard error of the work from all 10 sets for a given system was used for the error bars.

## Acknowledgements

This research is supported by National Science Foundation grant CHE 1945465. Simulations in this study have been performed primarily using Anton 2, Frontera, and Longhorn. We acknowledge COVID-19 HPC Consortium for providing access to these resources. This research is part of the Frontera computing project at the Texas Advanced Computing Center, made possible by National Science Foundation award OAC-1818253. Anton 2 computer time was provided by the Pittsburgh Super-computing Center (PSC) through Grant R01GM116961 from the National Institutes of Health. The Anton 2 machine at PSC was generously made available by D.E. Shaw Research. This research is also supported by the Arkansas High Performance Computing Center which is funded through multiple National Science Foundation grants and the Arkansas Economic Development Commission.

## Author Contributions

M.M. designed research; V.G.K. and D.S.O. performed simulations; V.G.K., D.S.O., U.H.I., A.P., J.L., and M.M. analyzed data and wrote the manuscript.

