## Supporting Information for "Differential Dynamic Behavior of Prefusion Spike Proteins of SARS Coronaviruses 1 and 2"

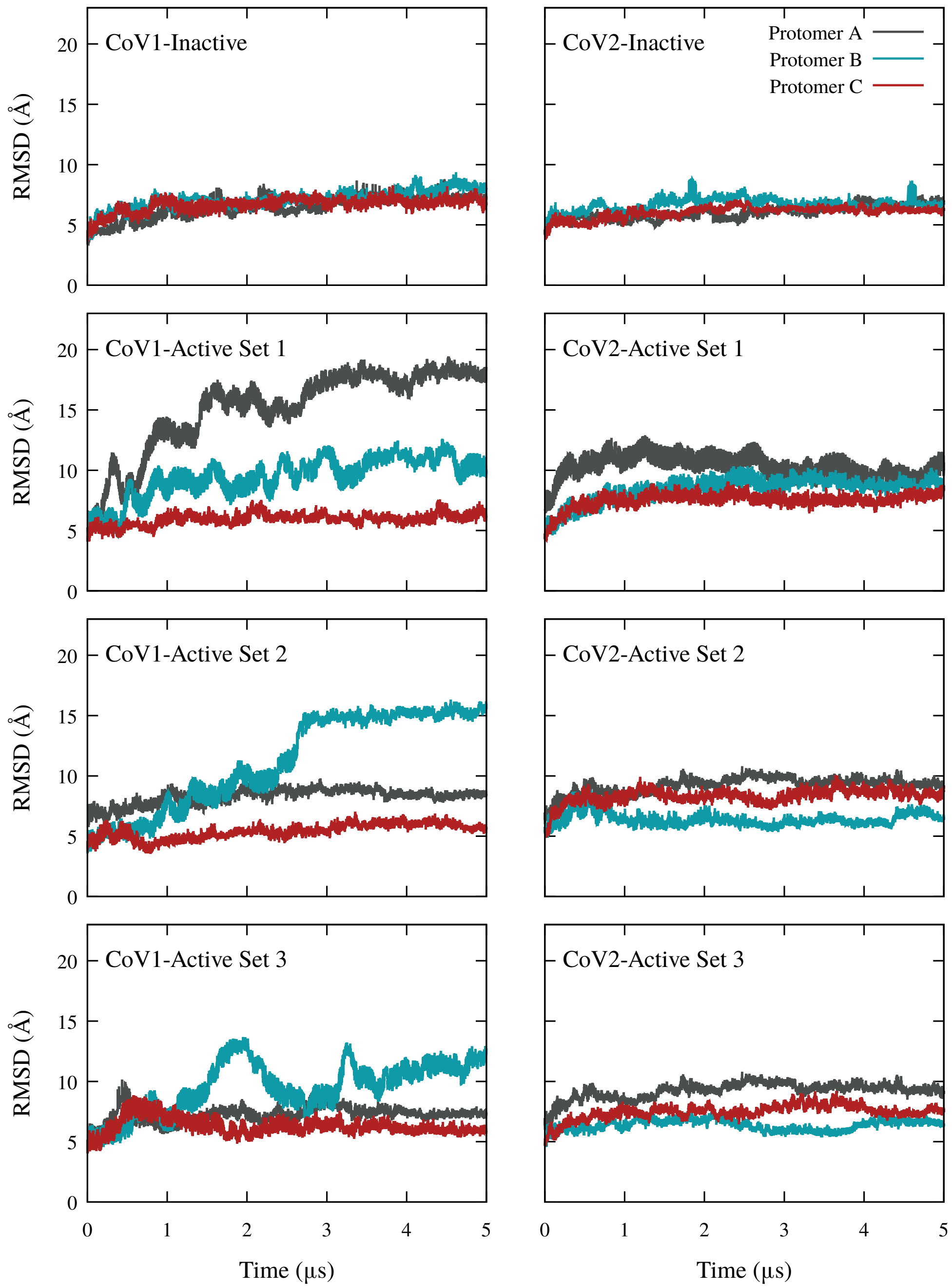


**Supplementary Figure 1: C-$\alpha$ RMSD for individual protomers.** The C-$\alpha$ RMSD calculated for each protomer relative to the initial cryo-EM structure over the 5 $\mu$s simulation is plotted for the inactive spike simulations and three sets of active spike simulations. Protomer A is colored dark grey, protomer B is colored light blue, and protomer C is colored dark red. The active CoV-2 spike protein is more stable overall than the active CoV-1 spike protein.

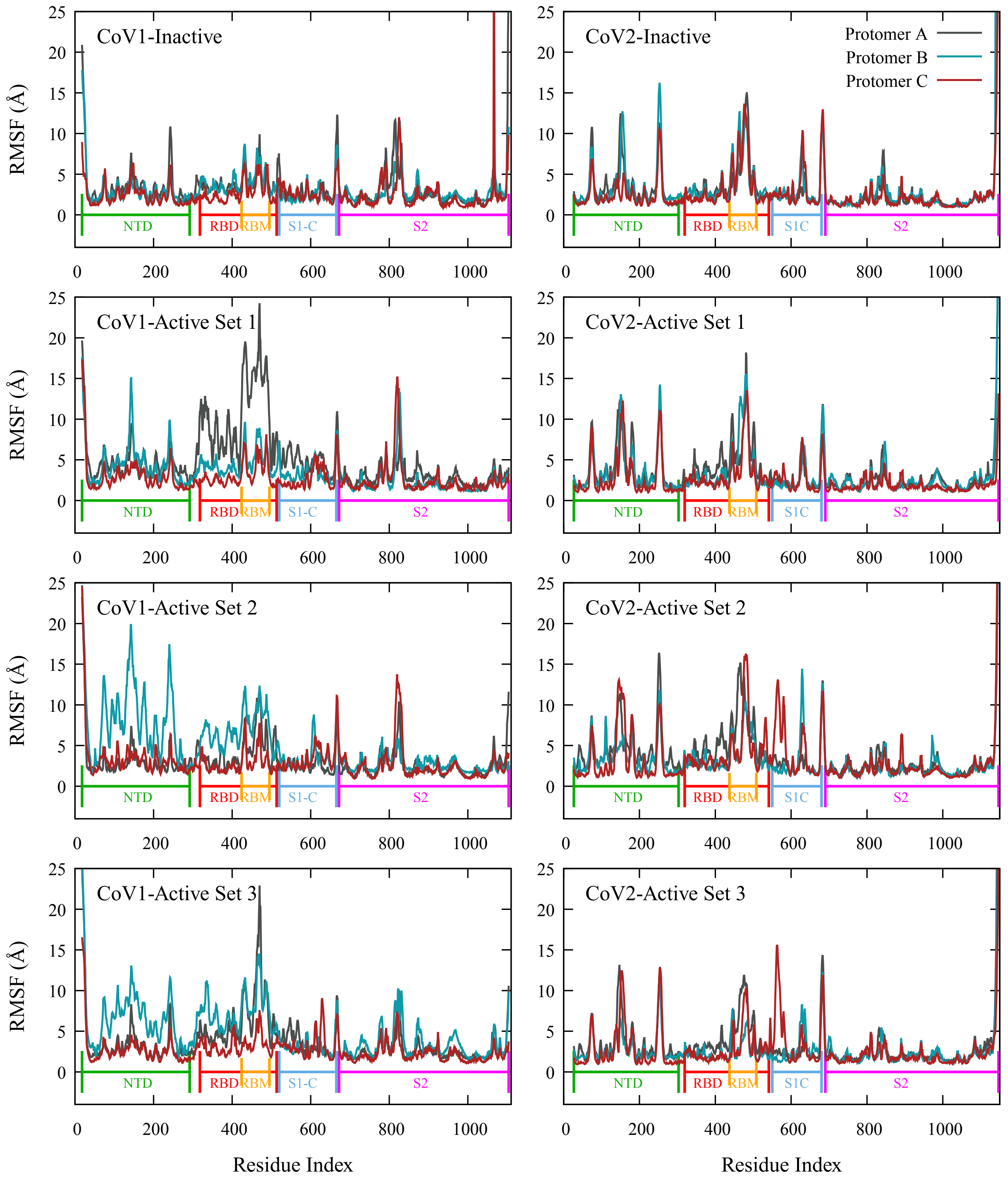


**Supplementary Figure 2: C-$\alpha$ RMSF for individual protomers.** The C-$\alpha$ RMSF for each protomer relative to the initial cryo-EM structure position was calculated for the inactive spike simulations and three sets of active spike simulations. Protomer A is colored dark grey, protomer B is colored light blue, and protomer C is colored dark red. The NTD and RBD of the active CoV-1 spike are more flexible than the corresponding regions of the active CoV-2 spike.

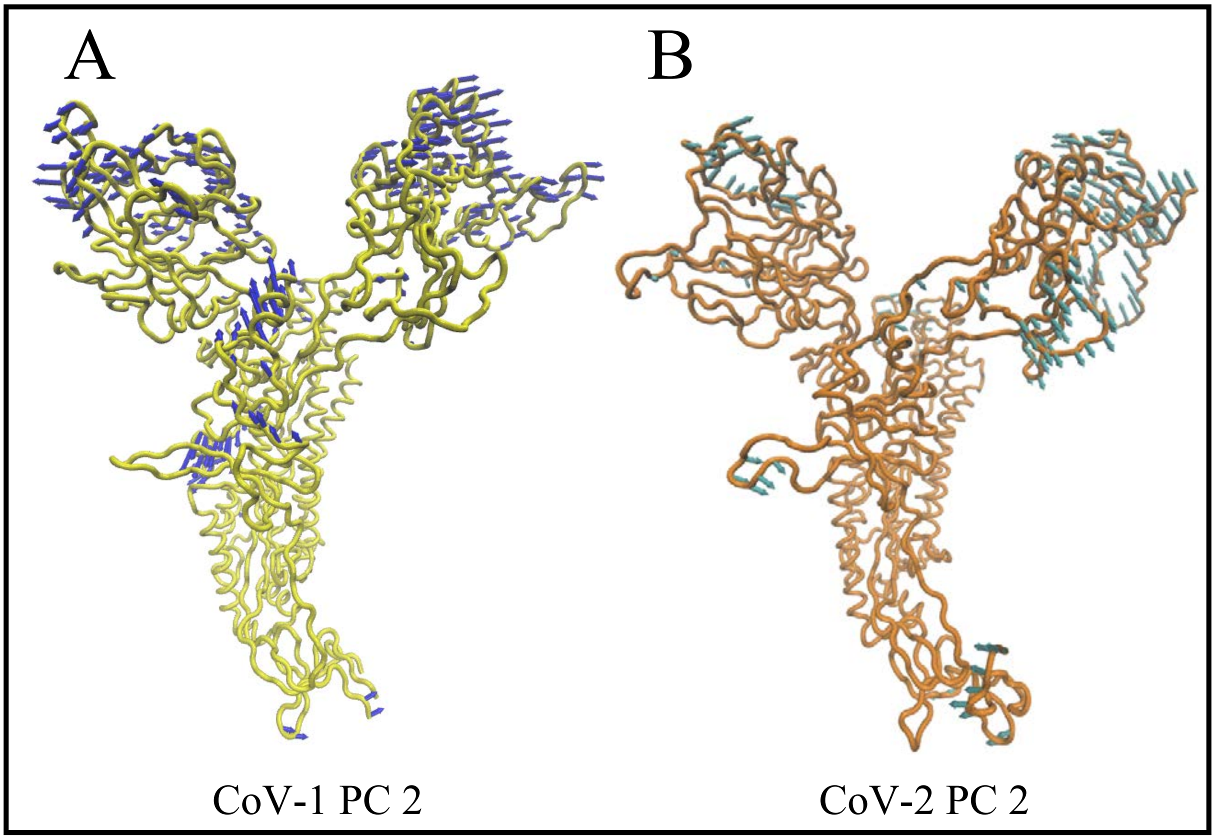


**Supplementary Figure** 3: **Visual representation of PC2 for all protomers in the inactive and active (Set 1) spike simulations for CoV-1 and CoV-2.** **(A)** Visual representation of PC2 for all Cov-1 protomers with the blue arrows at each C-$\alpha$ atom indicating direction and magnitude of variance. **(B)** Visual representation of PC2 for all CoV-2 protomers with the blue arrows at each C-$\alpha$ atom indicating direction and magnitude of variance. The NTD motions contribute more to the conformations sampled in the PC2 space than the PC1 space. These NTD motions are more pronounced in the CoV-1 spike, which also has more regions outside the NTD/RBD that show high variance.

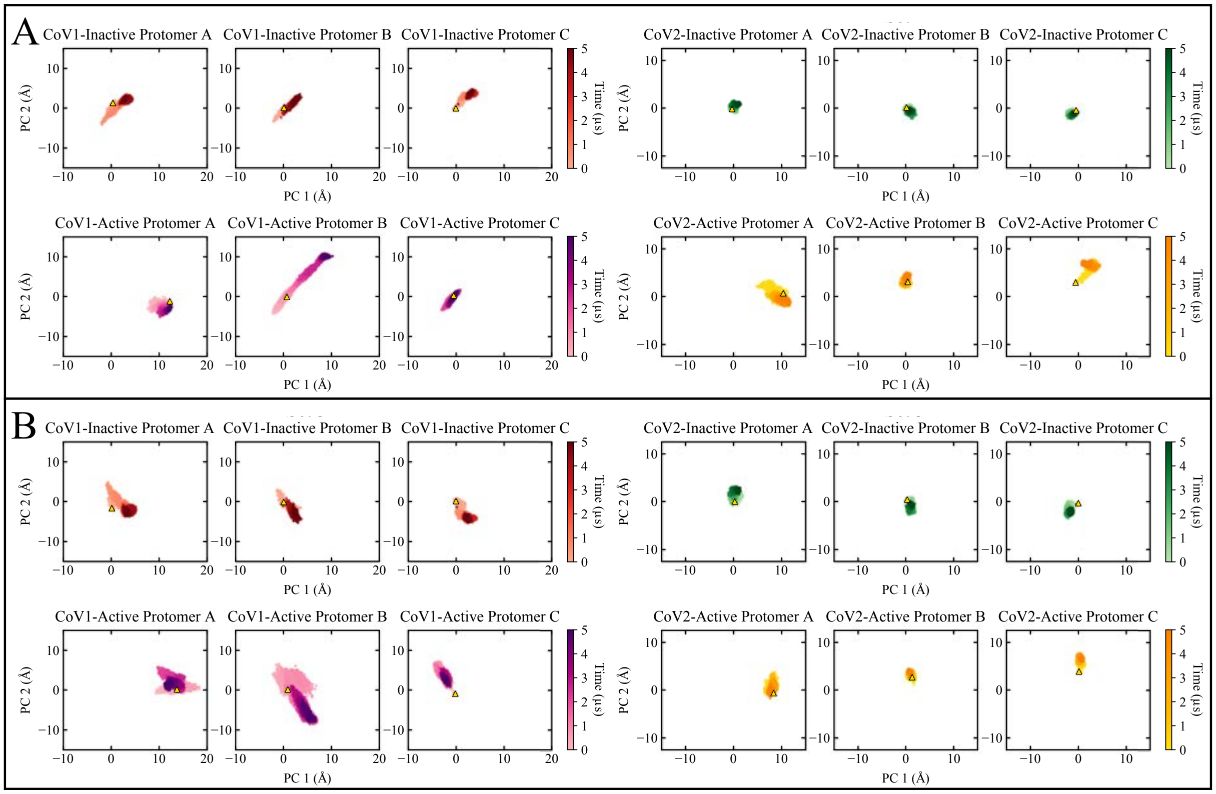


**Supplementary Figure 4: PCA of all protomers in the inactive and active (Sets 2 and 3) simulations for CoV-1 and CoV-2.** **(A)** Scatter plot of PC1 and PC2 for Set 2 of CoV-1 and CoV-2 active and inactive spike simulations.**(B)** Scatter plot of PC1 and PC2 for Set 3 of CoV-1 and CoV-2 active and inactive spike simulations. The coloring is the same as seen in Figure 3 with darker shades representing frames towards the end of the simulations. The active CoV-2 spike clearly samples fewer conformations in both PC1 and PC2 spaces than the active CoV-1 spike.

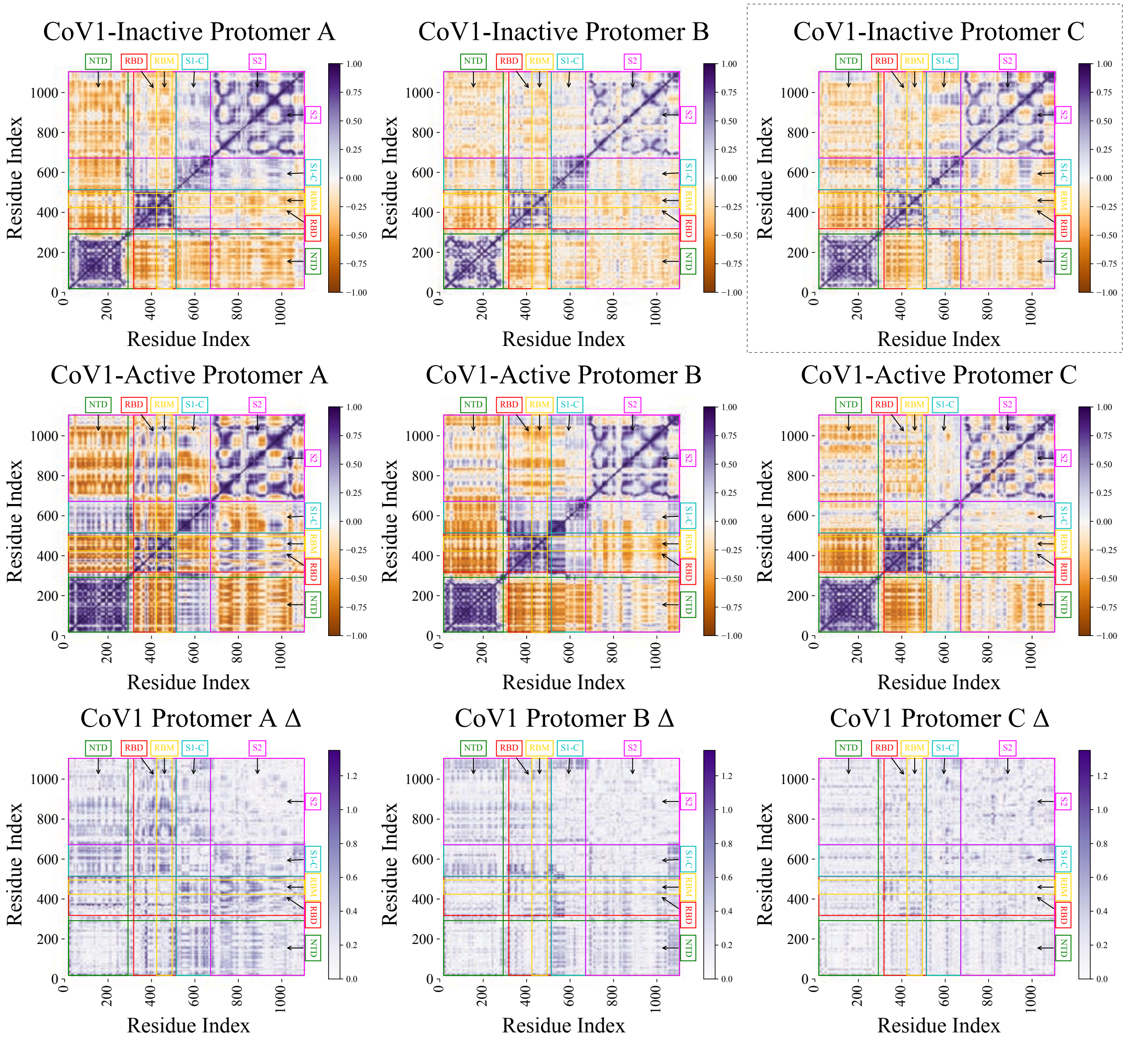


**Supplementary Figure 5: DNA correlation heat maps and Δ matrix for all protomers from the CoV-1 inactive and CoV-1 active (Set 1) spike simulations.** DNA heat maps showing the correlation of motions for the CoV-1 inactive protomers (first row), the CoV-1 active protomers from Set 1 (second row) and the difference matrices. The inactive protomer C correlation matrix, indicated by the dotted box, is the reference used for calculating the Δ matrix.

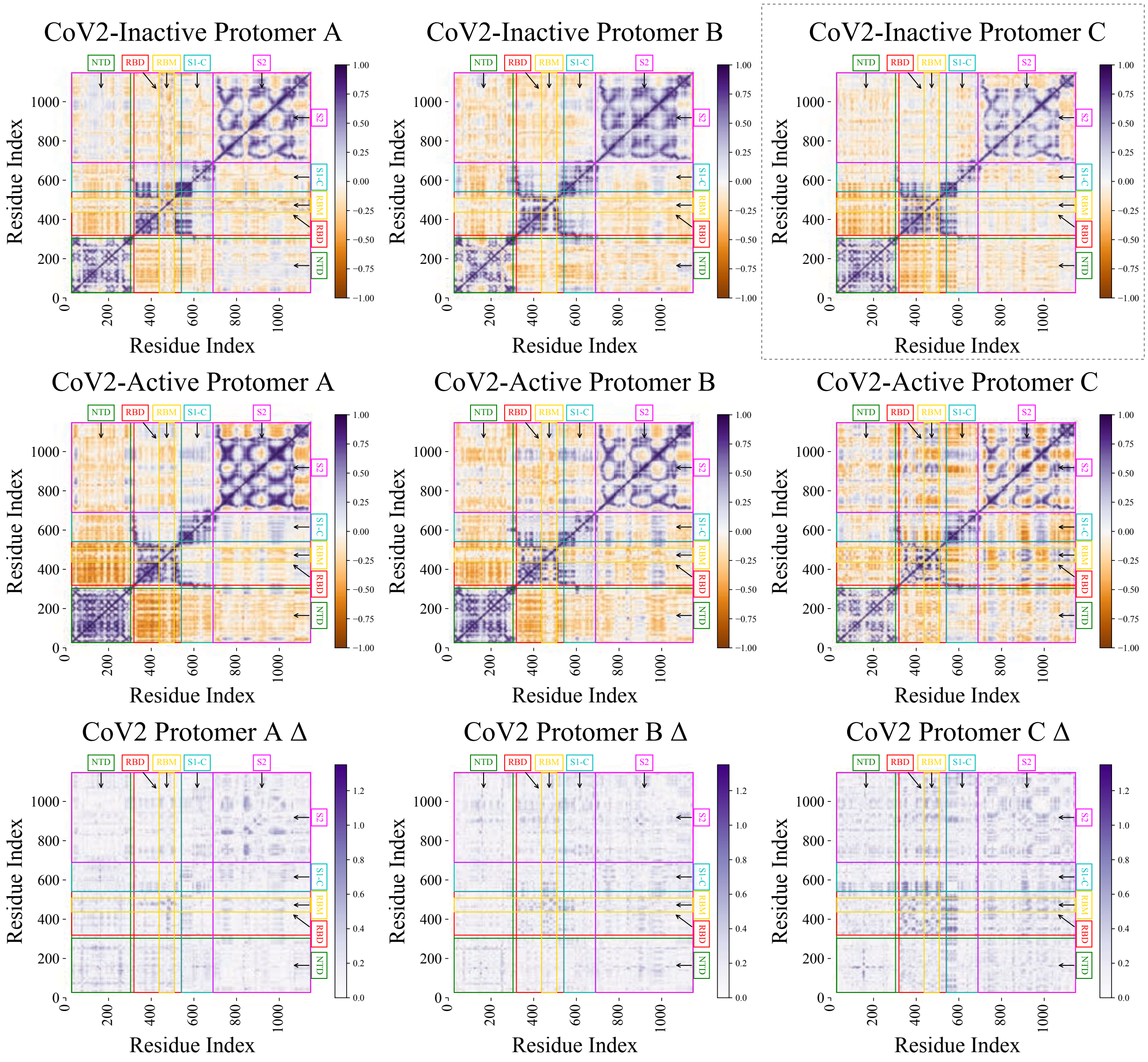


**Supplementary Figure 6: DNA correlation heat maps and Δ matrix for all protomers from the CoV-2 inactive and CoV-2 active (Set 1) spike simulations.** DNA heat maps showing the correlation of motions for the CoV-2 inactive protomers (first row), the CoV-2 active protomers from Set 1 (second row) and the difference matrices. The inactive protomer C correlation matrix, indicated by the dotted box, is the reference used for calculating the Δ matrix.

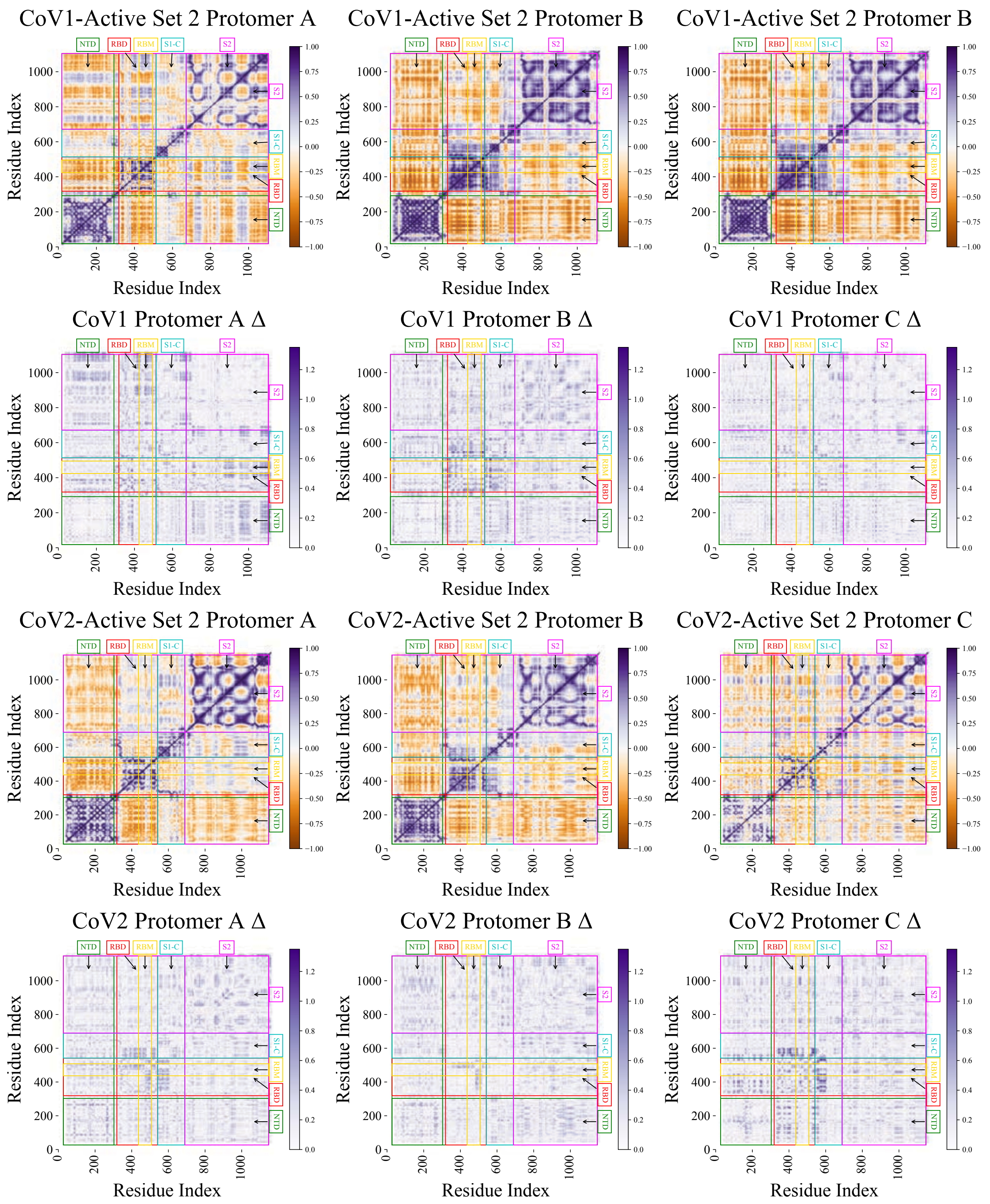


**Supplementary Figure 7: DNA correlation heat maps and Δ matrix for all protomers from Set 2 of the CoV-1 and CoV-2 active spike simulations.** **(A)** DNA heat maps showing the correlation of motions for the CoV-1 active (Set 2) protomers (first row) and the difference matrices (second row). The reference matrix from Figure S5 was used for Δ matrix calculations. **(B)** DNA heat maps showing the correlation of motions for the CoV-2 active (Set 2) protomers (third row) and the difference matrices (fourth row). The reference matrix from Figure S6 was used for Δ matrix calculations.

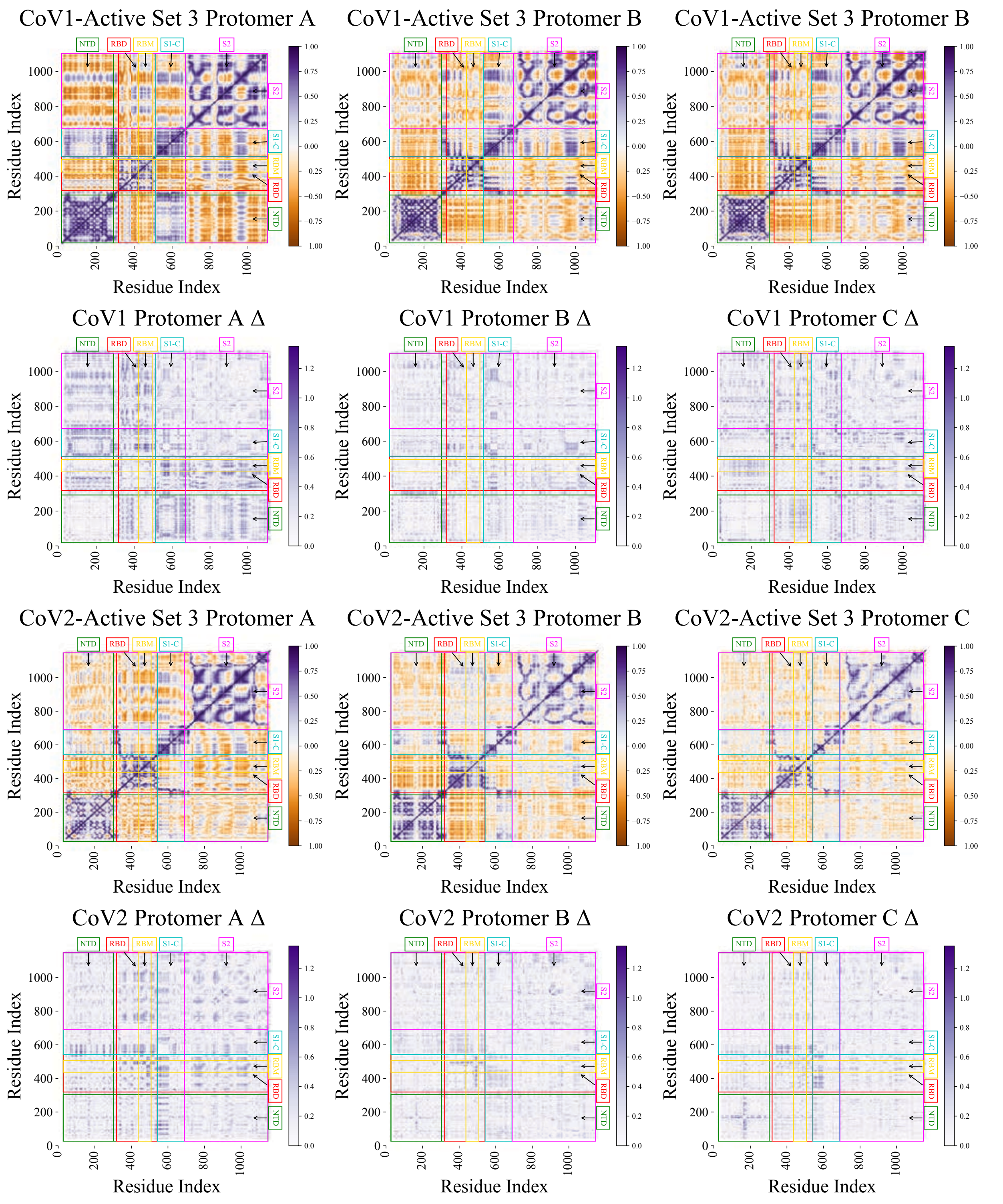


**Supplementary Figure 8: DNA correlation heat maps and Δ matrix for all protomers from Set 3 of the CoV-1 and CoV-2 active spike simulations.** **(A)** DNA heat maps showing the correlation of motions for the CoV-1 active (Set 3) protomers (first row) and the difference matrices (second row). The reference matrix from Figure S5 was used for Δ matrix calculations. **(B)** DNA heat maps showing the correlation of motions for the CoV-2 active (Set 3) protomers (third row) and the difference matrices (fourth row). The reference matrix from Figure S6 was used for Δ matrix calculations.

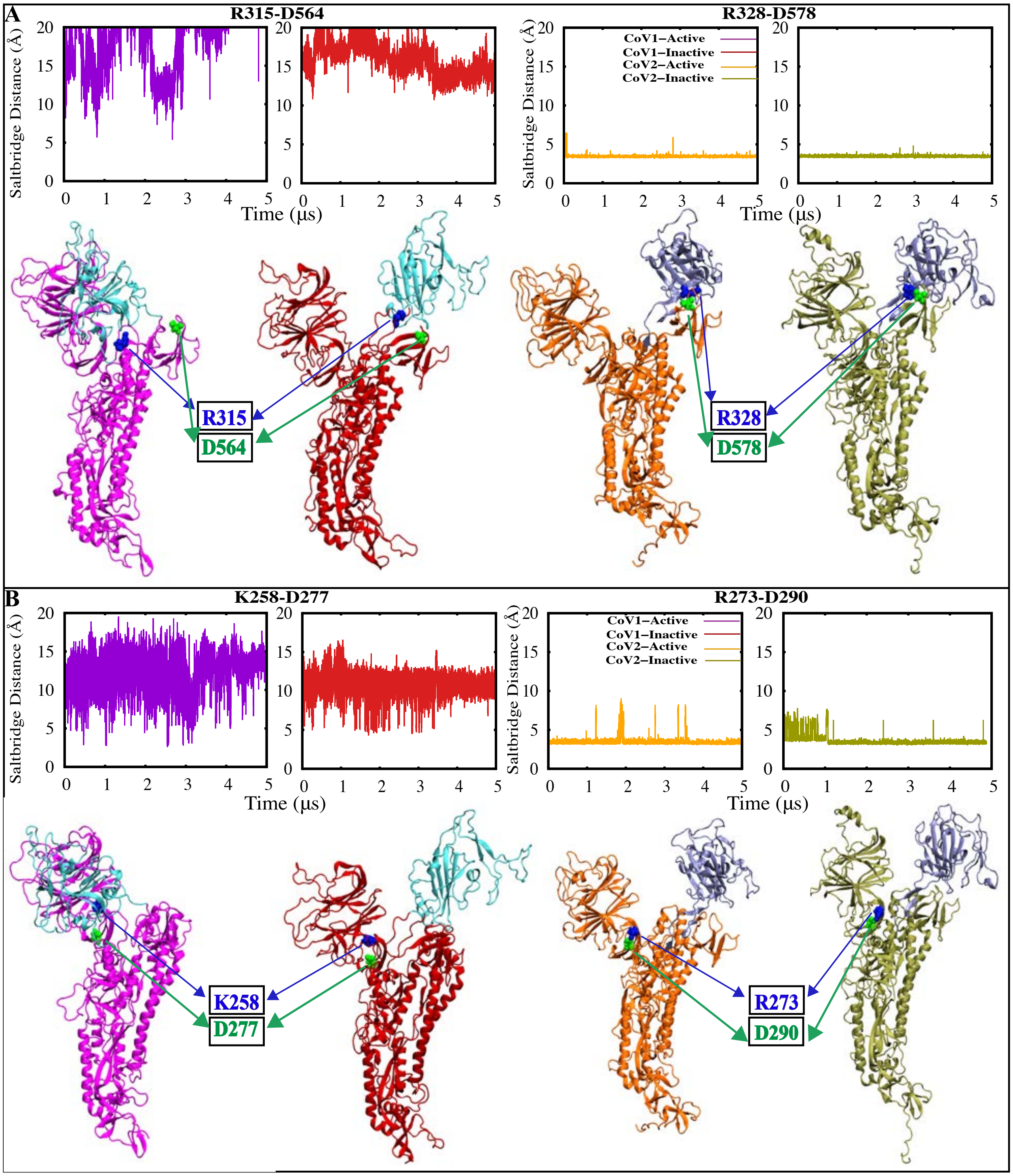


**Supplementary Figure 9: Conserved residues show distinct differential behavior in the CoV-1 and CoV-2 spike proteins.** Time series and visual representation of the minimum salt-bridge distance for **(A)** R315/328 (blue) - D564/578 (green) and **(B)** K258/R273 (blue) - D277/290 (green), shows that salt-bridges are formed in the CoV-2 spike protein but are absent in the CoV-1 spike protein. These salt-bridges potentially contribute to the higher relative stability of the CoV-2 spike protein. CoV-1 inactive is colored red, CoV-1 active is colored magenta, CoV-2 inactive is colored olive-green and CoV-2 active is colored orange.

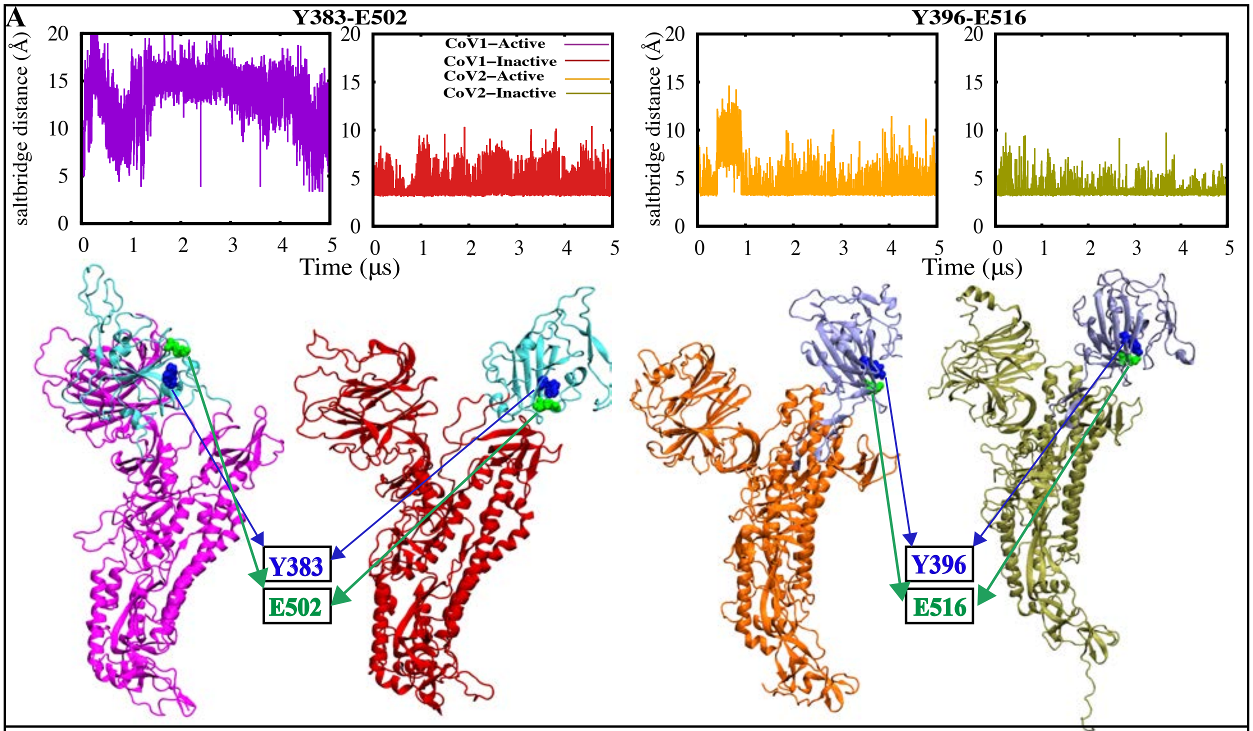


**B**

| | Occupancy (%)/Protomer | | |
|---|---|---|---|
| **R315-D564** | **A** | **B** | **C** |
| CoV1-active (set 1) | 0 | 0 | 0 |
| CoV1-active (set 2) | 0 | 0 | 0 |
| CoV1-active (set 3) | 0 | 0 | 0 |
| CoV1-inactive | 0 | 0 | 0 |
| **R328-D578** | | | |
| CoV2-active (set 1) | 100 | 76 | 0.31 |
| CoV2-active (set 2) | 59 | 11 | 0.95 |
| CoV2-active (set 3) | 98 | 5.2 | 0.16 |
| CoV2-inactive | 100 | 100 | 100 |

| | Occupancy (%)/Protomer | | |
|---|---|---|---|
| **K258-D277** | **A** | **B** | **C** |
| CoV1-active (set 1) | 0.96 | 2.4 | 1.9 |
| CoV1-active (set 2) | 0.34 | 0.12 | 0.45 |
| CoV1-active (set 3) | 0.09 | 0.34 | 0.61 |
| CoV1-inactive | 0.38 | 2.02 | 20 |
| **R273-D290** | | | |
| CoV2-active (set 1) | 98 | 64 | 94 |
| CoV2-active (set 2) | 100 | 98 | 85 |
| CoV2-active (set 3) | 100 | 96 | 96 |
| CoV2-inactive | 94 | 95 | 87 |

| | Occupancy (%)/Protomer | | |
|---|---|---|---|
| **Y383-E502** | **A** | **B** | **C** |
| CoV1-active (set 1) | 0.02 | 29 | 55 |
| CoV1-active (set 2) | 0.01 | 26 | 62 |
| CoV1-active (set 3) | 0.09 | 31 | 44 |
| CoV1-inactive | 64 | 21 | 98 |
| **Y396-E516** | | | |
| CoV2-active (set 1) | 65 | 66 | 94 |
| CoV2-active (set 2) | 27 | 67 | 80 |
| CoV2-active (set 3) | 31 | 26 | 45 |
| CoV2-inactive | 14 | 45 | 1.2 |

**Supplementary Figure 10: Hydrogen bond analysis for a conserved residue pair within the RBD. (A)** Time series and visual representation of the minimum H-bond donor-acceptor distance between Y383/396 (blue) and E502/516 (green), in the CoV-1 and CoV-2 spike respectively. CoV-1 inactive is colored red, CoV-1 active is colored magenta, CoV-2 inactive is colored olive green and CoV-2 active is colored orange. Table **(B)** shows the occupancy (%) of the salt-bridge and hydrogen-bond interactions between conserved residue pairs, for all protomers from all simulation sets.

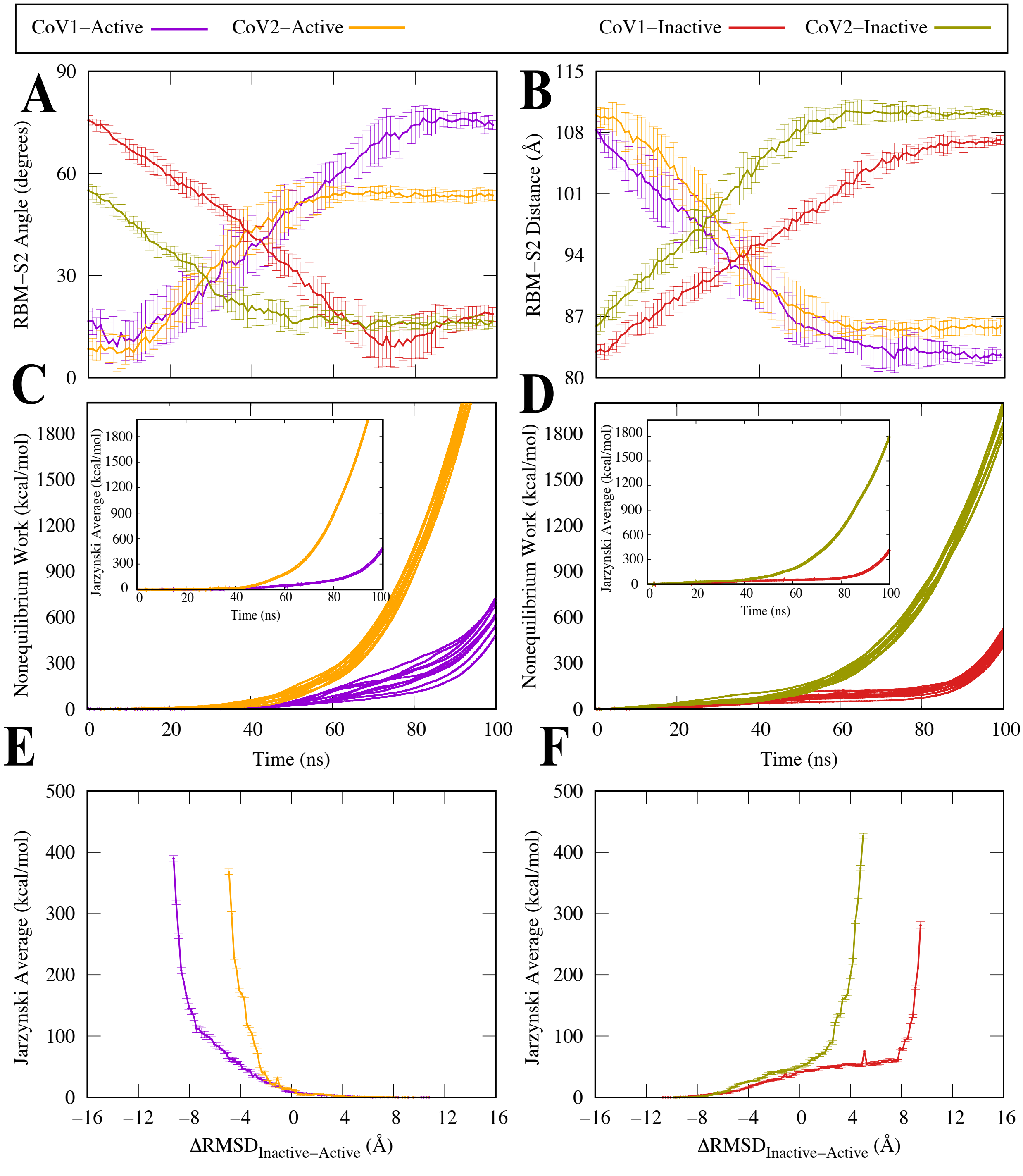


**Supplementary Figure 11: Three protomer SMD simulations. (A)** RBM-S2 Angle between the beta sheet region of the RBM and the alpha helical region of S2, shown as a function of time. Protomer activation is characterized by a decrease in the RBM-S2 angle. **(B)** RBM-S2 COM Distance between the beta sheet region of the RBM and the alpha helical region of S2, shown as shown as a function of time. Protomer activation is characterized by an increase in the RBM-S2 distance.**(C,D)** Accumulated non-equilibrium work as a function of simulation time. **(E,F)** Jarzynski average taken with respect to the $\Delta RMSD = RMSD_{Inactive}$ - $RMSD_{Active}$.
